# Decoupling the effects of nutrition, age and behavioral caste on honey bee physiology and immunity

**DOI:** 10.1101/667931

**Authors:** Miguel Corona, Belen Branchiccela, Shayne Madella, Yanping Chen, Jay Evans

**Affiliations:** Bee Research Laboratory, United States Department of Agriculture, Beltsville MD USA; Departamento de Microbiologia. Instituto de Investigaciones Biologicas Clemente Estable, Montevideo Uruguay

**Keywords:** Apis mellifera, nutritional stress, pollen, colony losses, division of labor, deformed wing virus, vitellogenin

## Abstract

Nutritional stress, and especially a dearth of pollen, is considered an important factor associated with honey bee colony losses. We used pollen-restricted colonies as a model to study the nutritional stress conditions experienced in colonies within intensively cultivated agricultural areas. This model was complemented by the establishment of an experimental design, which allowed us to uncouple the effect of nutrition, behavior and age in colonies of similar size and demography. We used this system to determine the effect of pollen restriction on workers’ behavioral development. Then, we analyzed the effect of nutritional stress, behavior and age on the expression of key physiological genes involved in the regulation of division of labor. Finally, we analyzed the effects of these variables on the expression of immune genes and the titers of honey bee viruses. Our results show that pollen restriction led to an increased number of precocious foragers and this behavioral transition was associated with important changes in the expression of nutritionally regulated physiological genes, immunity and viral titers. *Vitellogenin (vg)* and *major royal jelly protein* 1 (*mrjp1)* were the most predictive markers of nutrition and behavior. The expression of immune genes was primarily affected by behavior, with higher levels in foragers. Deformed wing virus (DWV) titers were significantly affected by behavior and nutritional status, with higher titer in foragers and increased levels associated with pollen ingestion. Correlation analyses support the predominant effect of behavior on immunity and susceptibility to viral infection, revealing that both immune genes and DWV exhibited strong negative correlations with genes associated with nursing, but positive correlations with genes associated with foraging. Our results provide valuable insights into the physiological mechanisms by which nutritional stress induce precocious foraging and increased susceptibility to viral infections.

## Introduction

Approximately 35-40% of the world’s crop production comes from plant species that depend on animal pollination [1] which is carried out primarily by honey bees [2]. However, populations of honey bees have experienced a severe decline during recent years worldwide [3, 4]. Possible causes for colony losses have been proposed, including the effects of pesticides [5], nutritional stress [6–8], the parasitic mite *Varroa destructor* [9, 10] and synergistic interactions between Varroa and honey bee viruses [11]. However, none of these independent factors is consistently associated with colony losses to suggest a single causal agent [3]. Thus, colony losses are likely the result of the combination of several underlying factors [12, 13].

There are several lines of evidence indicating that nutritional stress is an important contributing factor for colony losses. First, there is a positive relationship between the area of uncultivated forage land surrounding an apiary and annual colony survival [6, 8] suggesting that habitat loss play a significant role in honeybee colony losses. Second, at individual levels, honey bees from colonies kept in areas of high cultivation exhibited decreased levels of physiological biomarkers associated with nutrition compared with bees kept in areas of low cultivation [7, 14], demonstrating that availability of floral resources affects the nutritional state of honey bees.

Habitat loss associated with increased use of monocultures reduces both the quality (diversity of plant sources) and the quantity of the pollen collected by the bees [15]. Simultaneous flowering in monocultures provide pollen for a restricted period, and hives surrounded by monocultures presumably experience pollen dearth before and after this period. Pollen is the main source of proteins and lipids for honey bees [16, 17] and pollen from different plants differ in amino acid content [18, 19] and fatty acids [20]. Thus, the consumption of pollen from diverse plant sources increases the probability of obtaining the set of nutritional components (e.g., essential amino acids and fatty acids) required for balanced nutrition.

Plasticity is a key feature of honey bee behavioral development [21], which is important for understanding the consequences of nutritional stress on colony health and fitness. In the honey bee, as in other eusocial insects, behavioral division of labor in workers is initially age-dependent [21, 22]. Under typical spring and summer conditions in temperate climates, honey bee lifespan is approximately one month. During the first two to three weeks of adult life, workers perform tasks in the hive such as brood care (“nursing”); while in the last one to two weeks of life, workers forage for food outside the colony [23, 24]. However, honey bee behavioral development is flexible and can be accelerated or delayed according to colony demography and as an adaptive response to different challenges [21]. In particular, the timing of nurse to forager transition is normally accelerated in response to diverse types of stressors, including diseases [25] and malnutrition [26].

Interactions among key elements of the extended insulin/insulin-like growth signaling/target of rapamycin (IIS/TOR) pathway including vitellogenin (Vg), juvenile hormone (JH) and insulin-like peptides (ILPs), regulate important physiological processes in honey bees, including behavioral division of labor [27–31]. Vg is a yolk protein highly expressed in the fat bodies [29], secreted into the hemolymph and then imported by developing oocytes [32]. JH, one of the major insect hormones, is synthesized in the corpora allata in a final reaction catabolized by the enzyme methyl farnesoate epoxidase (MFE) [33]. There are two genes encoding insulin genes (*ilp1* and *ilp2*) in the honey bee genome [34], which are expressed in several tissues and organs including the brain and fat bodies [27, 29, 30]. In workers, negative interactions among *vg* with JH and insulin like peptides are associated with task performance: while *vg* levels are higher in nurses compared with foragers [29, 35–37], JH titers [21, 36, 38, 39] brain *ilp1* and fat body *ilp2* mRNA levels follow an opposite pattern [27, 29, 30]. Experimental manipulations showed that these correlations reveal causative positive and negative interactions among these three regulatory elements. Thus, JH treatments inhibit *vg* expression [35, 40], but induce higher brain *ilp1* levels [29] and precocious foraging [21, 41]. Consistently, experimental reduction of *vg* by RNA interference also result in higher levels of JH [27] and precocious foraging [28].

In recent years, the proposal that nutrition is an important factor regulating the division of labor among social insects has received increased support [26, 30, 42–46]. Pollen consumption is low in young bees, largest in about 9-day-old nurses and then declines to minimal amounts in foragers [47], which rely primarily on honey intake. Cage experiments showed that 6-day-old workers eating pollen and sugar have higher abdominal *vg* and low brain *ilp1* levels, while same-age bees fed with only sugar had the inverse pattern [30]; resembling the expression profiles observed in colony-reared nurses and foragers, respectively [29]. The effect of pollen on behavioral development could result from its protein content. In support of this view, it has been shown that increased levels of hemolymph amino acids, induced by protein injection, resulted in high vg mRNA levels and delayed foraging [27]. Additionally, lipid metabolism also has been involved in nutritional regulation of division of labor. Lipid content closely correlates with task performance, being higher in nurses compared to foragers [43]. Feeding of an inhibitor of fatty acid synthesis (TOFA) induced precocious foraging [48], further supporting a role of lipids on the regulation of division of labor. However, it remains to be tested whether supplementary lipids (e.g., essential fatty acids) have an effect on the nutritional regulation of this process.

Since genes that regulate behavioral division of labor are nutritionally regulated, they could be expected to be ideal biomarkers of both nutritional and behavioral state in cage as well as colony-level studies. Progressive evidence is revealing that behavioral and nutritional states influence honey bee immunity and susceptibility to pathogens. Thus, the use of reliable markers for these traits can also facilitate effective disease diagnosis. Indeed, the initial use of *vg* as a predictive marker of colony losses [49] and colony nutritional state [14] provides a promising starting point toward this direction.

Although insects lack an acquired immunity, they have developed an efficient innate immune system against a broad variety of pathogens [50]. Innate immunity has been traditionally divided into humoral and cellular responses, but there are complex interactions and no clearly defined boundaries between both types of immune responses [51]. Four main gene pathways mediate insect innate immunity: Toll, IMD, JNK and JAK/ STAT, which include elements implicated in both the humoral and cellular immune responses and belong to gene families involved in signaling, pathogen recognition and effector functions [52, 53].

Humoral immune responses are mainly associated with antimicrobial effectors belonging to the Toll pathway such as *abaecin* [54], *defensin1 (def1)* [55] and *hymenoptaecin* (*hym*) [56]. On the other hand, cellular immunity is mediated by haemocytes and involves responses including phagocytosis, nodulation and encapsulation [57] While phagocytosis is mediated by pathogen recognition proteins (e.g., peptidoglycan recognition proteins L and S) [58] and transmembrane proteins such as *eater* [59], nodulation and encapsulation are often accompanied by melanization, a process catalyzed by the pro-phenoloxidase (*PPO*) and pro–phenoloxidase activator (PPOact) enzymes [52, 60, 61].

Different studies have revealed interactions among nutrition, immunity and physiological changes associated with behavioral development. First, nutrition affects the expression of immune genes [62–64] and susceptibility to different pathogens [65–68]. Second, infections of pathogens and parasites, such as the microsporidia *Nosema ceranae*, are associated with high JH and low *vg* levels and precocious foraging [25]. However, little is known about the molecular mechanism by which colony nutritional conditions lead to changes in behavioral development, susceptibility to pathogens and ultimately, colony survival.

In this study, we used pollen-restricted colonies as a model to study the nutritional stress conditions occurring in colonies within intensively cultivated agricultural areas. We used a new technique for colony establishment, which allow us to uncouple the effects of age, behavior and nutrition. We first analyzed the effect of pollen restriction on foraging and brood production and then we used qPCR to analyze the role of pollen restriction on molecular markers of nutrition, immune gene expression and dynamic of viral infection. Finally, we used a multidimensional analysis to show that nutrition has a major role in the dynamics of key genes associated with behavior, nutrition and immunity. This study opens new avenues in the understanding of the mechanisms underlying the effects of nutrition on behavioral development, immunity and susceptibility to virus infections. The results obtained also have important implications for the understanding of the consequences of the pollen deprivation on the induction of worker’s abnormal accelerated behavioral development and reduction in colony population associated with colony collapse.

## Results

### 1. Effect of pollen restriction on foraging

Both groups of pollen-restricted (P-R) and pollen-unrestricted (P-UR) bees started foraging when they were 11 days old. Observations were continued until the bees were 20 days old. During the total observation period, the average number of new foragers observed in the P-R colonies was almost 3X higher when compared with P-UR colonies (100 and 35, respectively; P= <0.0001). Analysis of nectar and pollen foragers in each nutritional group showed higher numbers of both types of foragers in P-R compared with P-UR colonies (nectar: P-R=53, P-UR-24, P= <0.0001; pollen: P-R=47, P-UR=11, P= <0.0001). While no significant differences were found between the number of nectar and pollen foragers in P-R colonies (P=0.52), the difference between both types of foragers in P-UR colonies was highly significant (P=0.0027) with a lower number of new pollen foragers than nectar foragers under this condition (Figure 1A and Figures S2 and S3).

**Figure 1.**
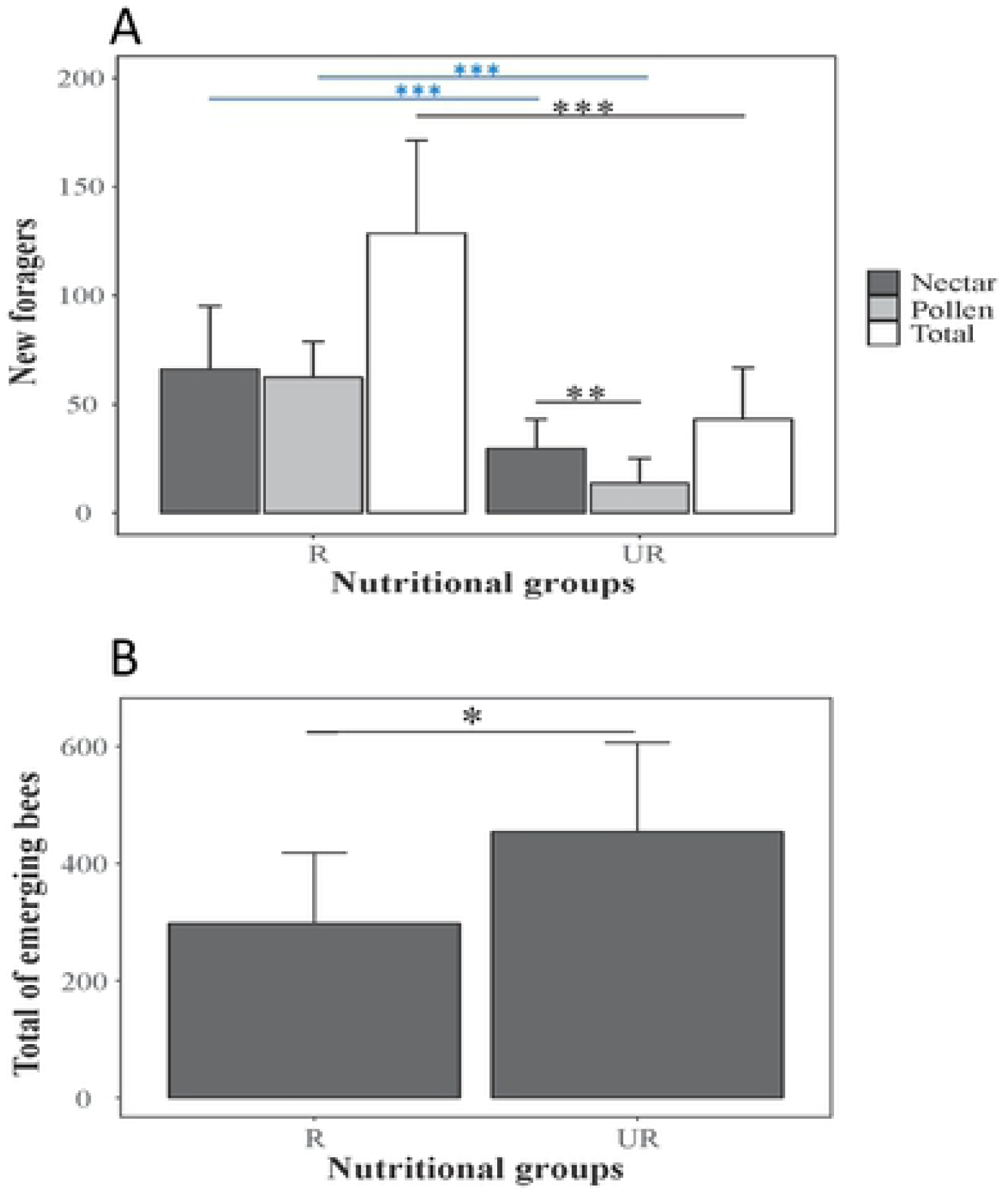
Effects of pollen restriction on foraging and brood production. Values were pooled from eight pollen restricted (P-R) and eight pollen un-restricted colonies (P-U). The upper panel (A) shows the effect of nutrition on the average number of new foragers per day. The y-axis indicates the number of nectar foragers, pollen foragers and total foragers observed per day (sum of 3 periods of 15 min each). Nutritional groups plotted in the x-axis are as follow: pollen-restricted colonies (R), pollen-unrestricted colonies (UR). The lower panel (B) compares the total number of bees born (y-axis) from colonies in both nutritional conditions (x-axis). Error bars represent SE across colonies. Significant differences in the total number of foragers between R and UR colonies are represented with black stars. Differences in pollen or nectar foragers between nutritional conditions are represented with blue stars.

### 2. Brood production

Two days after the end of the behavioral observations, and one day after the last collection of 3-week-old workers, newly-emerged bees were counted on frames from P-R and P-UR colonies until the last bees emerged after 16 days. The cumulative number of newly-emerged bees from P-R colonies was significantly lower compared to P-UR colonies (2573 and 3881, respectively. P=0.014) (Figure 1B).

### 3. Behavioral, nutritional and age analyses of gene expression and virus titers Genes associated with physiological activity

*Vg* expression was significantly affected by behavior, with higher levels in nurses in all nutritional and age groups (P= <0.0001 for all behavioral comparisons). Expression levels in nurses were on average 3.75X higher compared with foragers. (Figure 2A). Nutritional treatments also had a significant effect on *vg* expression, with higher levels in nurses from colonies with unrestricted access to pollen (P-UR) compared with nurses from colonies with restricted access to pollen (P-R) in both age groups (2-week-old: P= <0.0001; 3-week-old: P=0.0075). Foragers also showed higher levels of *vg* in P-UR colonies compared with P-R colonies, but only at the younger group (P=0.0003). Lastly, age comparisons showed a significant increase in 3-week-old workers in all the nutritional and behavioral comparisons with the exception of foragers from P-UR colonies (Figure 2A and Table S2).

**Figure 2.**
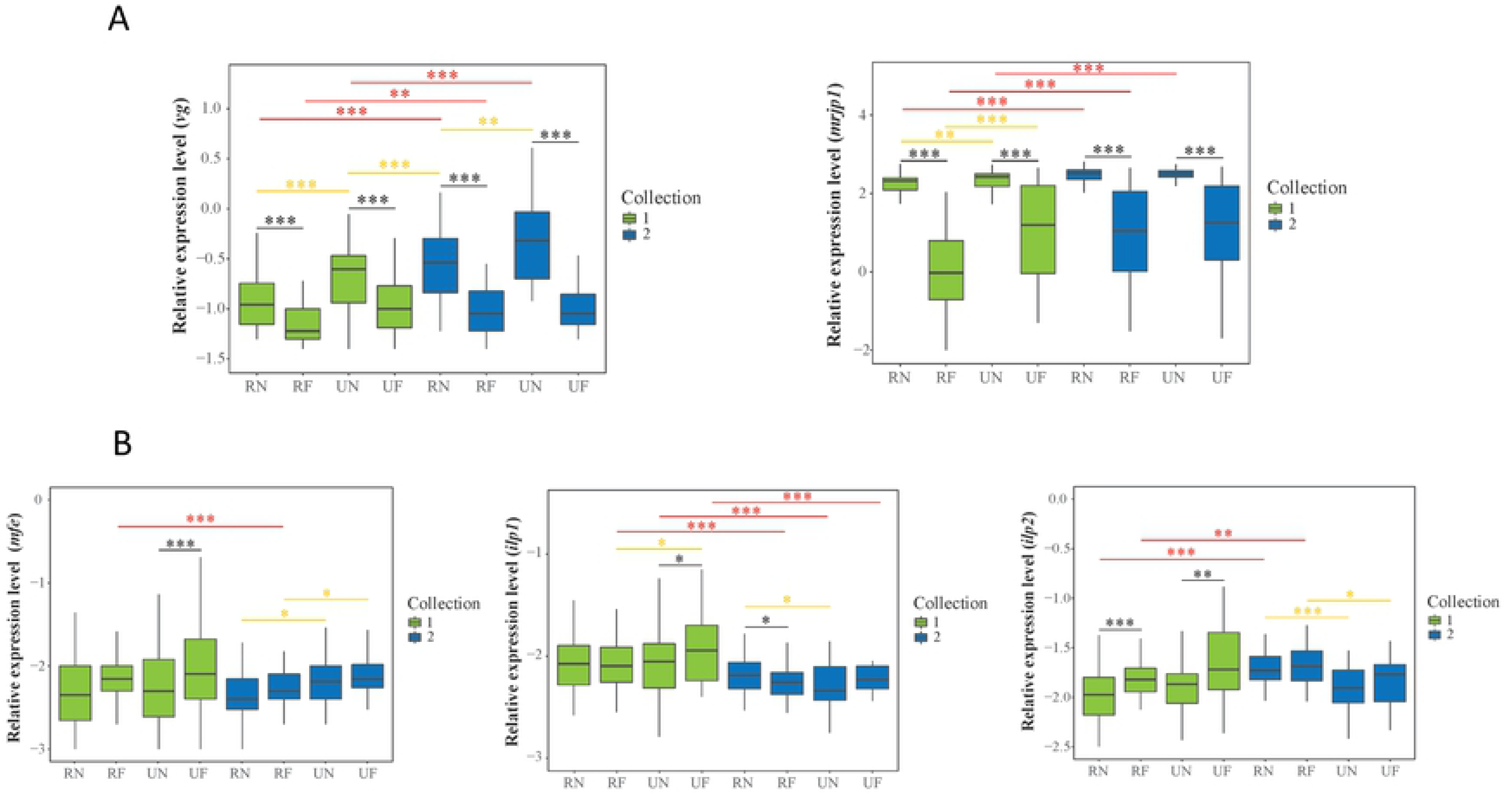
Relative mRNA levels of genes associated with physiological activity and behavior. The upper panel (A) compares the expression of genes associated with nursing: *vg* and *mrjp1*. The lower panel (B) compares the expression of genes predominantly associated with foraging: *mfe*, *ilp1* an *ilp2*. The y-axes indicate the relative expression levels relative to the ribosomal protein RPS5 (control) gene mRNA level. Boxes show the 1^st^ and 3^rd^ interquartile range and median are represented by a line. Whiskers include the values of 90% of the samples. Black, yellow and red stars represent significant differences between pairs of behavioral, nutritional and age groups, respectively.

*Mrjp1* expression levels were strongly affected by behavior with higher levels in nurses compared with foragers in all the nutritional and age comparisons. On average, *mrjp1* levels in nurses were 5X higher compared with foragers. On the other hand, nutritional comparisons showed that pollen consumption induced a significant increase of *mrjp1* in younger nurses (P=0.009) and foragers (7.2e-6) compared with workers from P-R colonies. No nutritional differences were observed in older 3-week-old workers. Age comparisons showed a tendency toward higher levels in older foragers, which become significant in all the comparisons with the exception of foragers from P-UR colonies. It is interesting to note, that our whole-body analyses reveal that *mrjp1* mRNA levels were substantially higher compared with *vg*, being on average over 500X higher (Figure 2A).

*Mfe* expression showed behaviorally related changes in 2-week-old workers, with higher levels in nurses compared with foragers (P-R= 0.015, P-UR=0.016). In contrast, no significant behavioral differences were observed in 3-week-old workers. On the other hand, nutritional comparisons showed significantly higher *mfe* expression associated with pollen intake exclusively in 3-week-old workers (nurses: P=0.035, foragers: P=0.014). Lastly, age comparisons revealed a trend toward down-regulation of *mfe* expression in older workers. However, significant differences were only found in foragers from P-R colonies (P=0.002). (Figure 2B).

The expression of *ilp1* revealed complex interactions in our behavioral and nutritional analyses. The behavioral analysis showed significantly higher levels in younger foragers from P-UR colonies (P=0.033) and in older nurses from P-R colonies (P=0.034). On the other hand, nutritional comparisons exhibited higher levels in 2-week-old foragers from P-UR colonies and 3-week-old nurses from P-R colonies (2-week-old: nurses P=0.61, foragers P=0.014; 3-week-old nurses P=0.01, foragers P=0.30). In contrast with behavior and nutritional analyses, age comparisons revealed a significant down-regulation in the expression of *ilp1* in most of the nutritional and behavioral groups of older workers (P-R nurses: P=0.058, P-R foragers: P= <0.0001, P-UR nurses: P= 0.0001, P-UR foragers: P= 0.0005) (Figure 2B).

Expression of *ilp2* showed behavioral differences only in younger workers, with higher levels in foragers both from P-R (P=0.0001) and P-UR (P=0.006) colonies. Nutritional comparisons showed a trend toward higher *ilp2* levels with pollen consumption in 2-week-old workers, but the differences were marginally significant (P-R=0.08, P-UR=0.08) (Table S2). In contrast, higher expression of *ilp2* was associated with pollen restriction in both 3-week-old nurses (P=0.0001) and foragers (P=0.037). Age comparisons revealed higher *ilp2* expression in workers from colonies experiencing pollen restriction (nurses P= <0.0001, foragers P=0.001). Additionally, our qPCR analyses revealed, that while *Ilp1* and *mfe* are expressed at similar levels, *ilp2* is expressed on average 2X higher compared with these genes (Figure 2B).

#### Immune genes

We analyzed the expression of 9 representative immune genes: *dorsal*, *def1, abaecin*, *hym*, *pgrp-lc*, *pgrp-sc*, *eater*, *ppo-act* and *ppo* (36 gene/age/nutritional/behavioral comparisons) (Figure 3 and 4 and Table S2). Overall behavioral analyses of these immune genes showed a strong trend toward higher levels in foragers, reaching significant differences in 30 out of 36 comparisons (83.3%). Remarkably, none of the comparisons showed higher expression levels of these genes in nurses than in foragers. On the other hand, nutrition had a significant effect on the expression of immune genes in 12 out 36 comparisons (33.3%). A higher percentage of comparisons showed up-regulation associated with pollen ingestion (22%) than showed up-regulation associated with pollen-restriction (11%). Lastly, age analysis of immune genes revealed significant differences in 20 out of 36 comparisons (55.5%). From these comparisons, 70% of them showed age-related increased expression and 30% age-related down-regulation.

**Figure 3.**
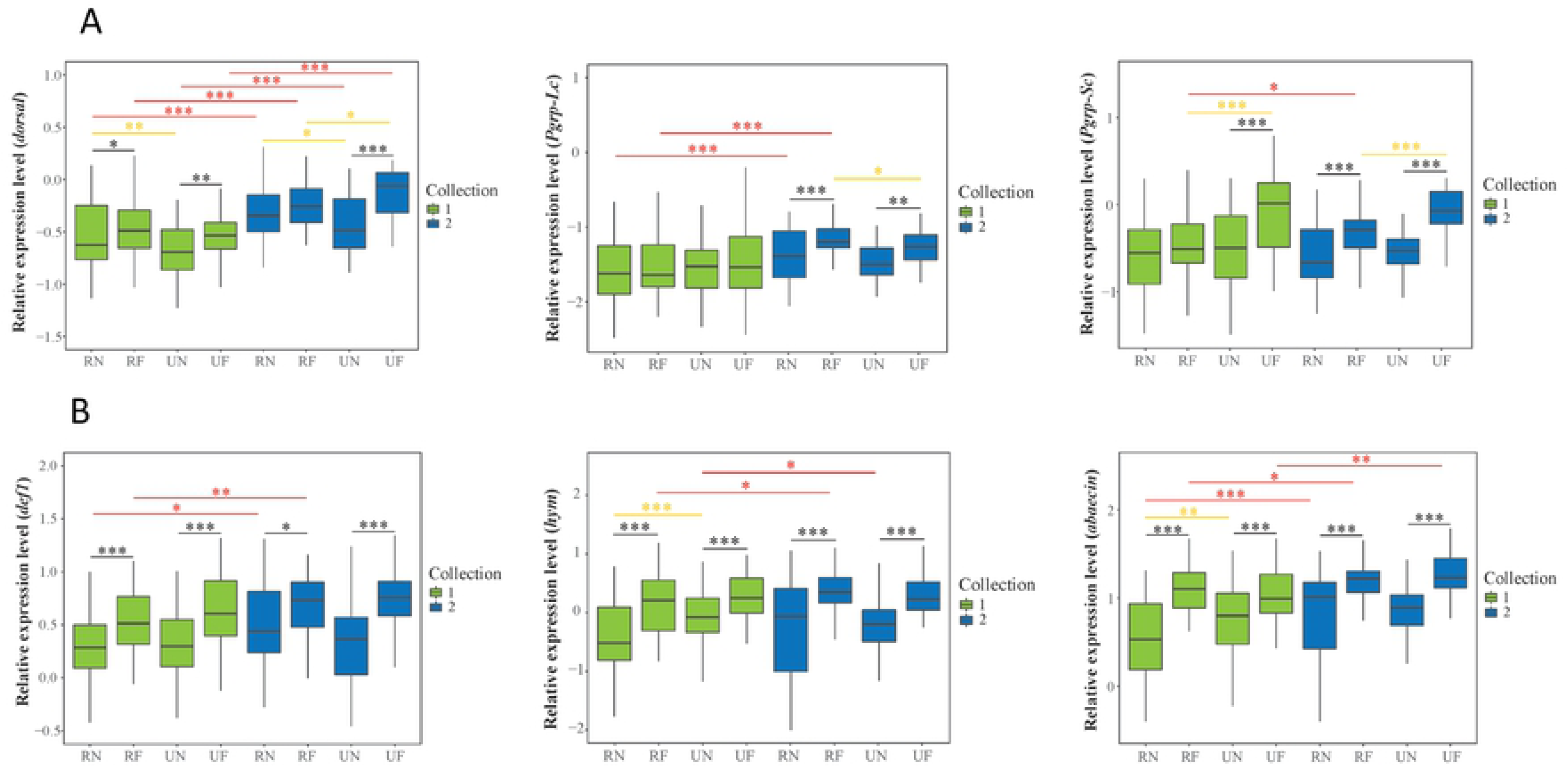
Expression of immune genes predominantly involved in the humoral immune response. The y-axes indicate the relative expression levels relative of *dorsal*, *def-1*, *abaecin*, *hym pgrp-lc* and *pgrl-sc* relative to the ribosomal protein RPR5 (control) gene mRNA level. In the gene which values showed normal distribution and homogeneity (*dorsal*), the boxes show media +/- standard deviation and the whiskers include the minimum and maximum values. In the graphs of the genes which values exhibited non-parametric distribution (*def-1*, *abaecin*, *hym pgrp-lc* and *pgrl-sc)* the boxes show the 1^st^ and 3^rd^ interquartile range and the whiskers include the values of 90% of the samples. In both cases a line represents the media. Significant differences between groups are represented as in figure 2.

**Figure 4.**
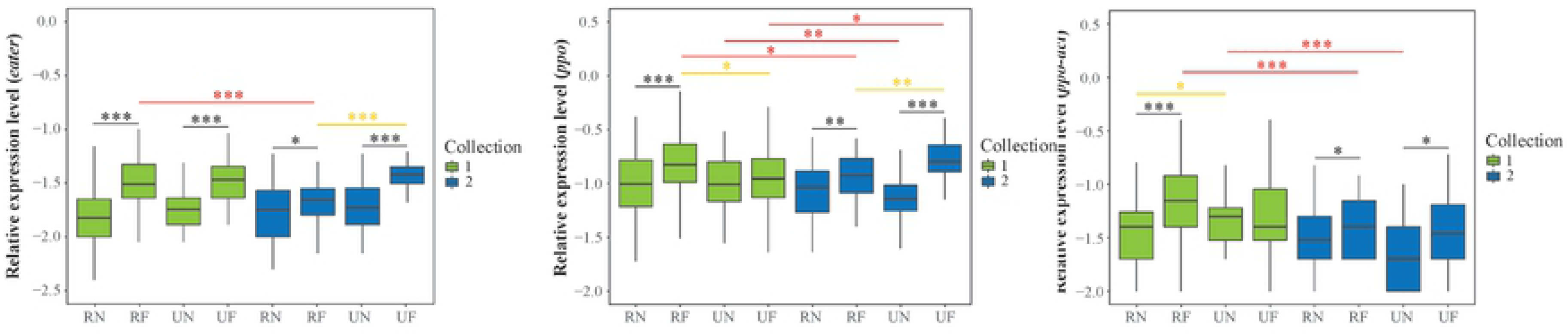
Expression of genes predominantly involved with the cellular immune response. The y-axes indicate the relative expression levels of *eater*, *ppo* and *ppo-act* relative to the ribosomal protein RPR5 (control) gene mRNA level. The boxes of the gene which expression values showed normal distribution (*ppo*) or the genes which values exhibited non-parametric distribution (*eater* and *ppoact*) are represented as in the Figure 3.

Gene-specific behavioral analyses revealed that the immune genes exhibiting the highest percentage of comparisons (100%) with higher levels in foragers include the three antimicrobial effectors analyzed (*def1*, *abaecin* and *hym)*. Conversely, the pathogen recognition protein *pgrp-sc* was the immune gene less affected by behavior (50% of the four comparisons). The immune gene most affected by nutrition was the regulatory gene *dorsal* (75%), while *def1* was not affected by nutrition. D*orsal* also was the immune gene most influenced by age with a significant effect in 100% of the comparisons, followed by *abaecin* and *ppo (75%)*. In contrast, *eater* and *pgrp-sc* were the immune genes less affected by age (25% of the comparisons) (Figure 3 and 4, Table S2).

#### Virus titers

We analyzed the effect of behavior, nutrition and age on the titers of three of the more prevalent honey bee viruses, namely deformed wing virus (DWV), black queen cell virus (BQCV) and sacbrood virus (SBV) in12 age/nutritional/behavioral comparisons (Figure 5, Table S2). Overall behavioral analysis of these viruses revealed significant differences in 50% of the comparisons. From these comparisons, most of them were upregulated in foragers (83.3%) compared with nurses (16.6%). Nutritional analysis showed significant differences only in 25% of the comparisons, all of them observed in younger workers.

**Figure 5.**
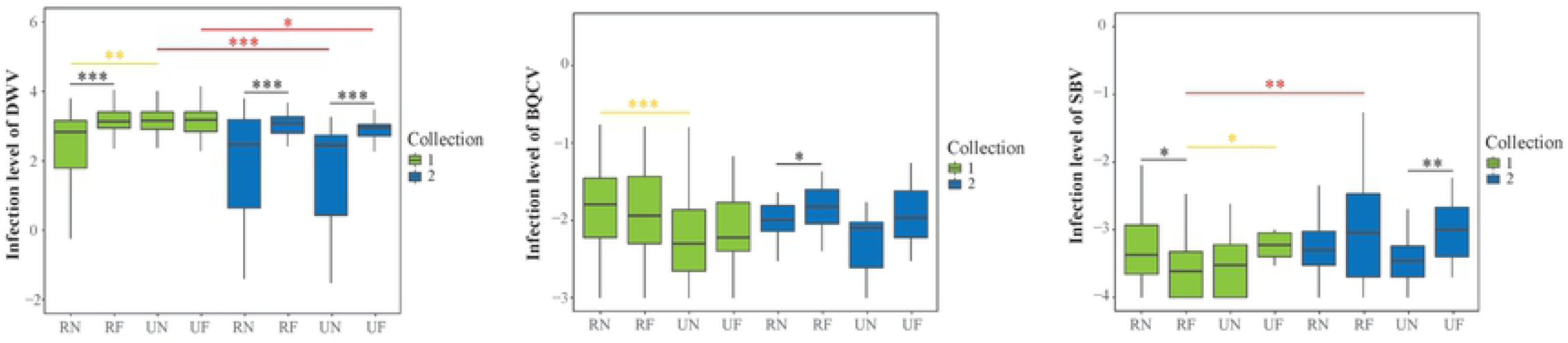
Relative levels of honey bee ARN viruses. The y-axes indicate the titers of DWV, SBV and BQCV relative to the ribosomal protein RPS5 (control) gene mRNA level. Boxes show 1^st^ and 3^rd^ interquartile range and the median is represented by a line. Whiskers include the values of 90% of the samples.

Virus-specific behavioral analyses showed that DWV had a strong trend toward higher titers in foragers, with highly significant differences in all nutritional and age groups (P= <0.0001) with the exception of young foragers from P-UR colonies. Nutritional comparisons revealed higher titers of DWV only in younger nurses from P-UR colonies (P=0.0001), while no significant differences were observed in other comparisons among nutritional, behavioral and age groups. On the other hand, age comparisons showed a tendency toward lower levels in older workers, with significant differences in nurses (P=<0.0001) and foragers (P=0.015) from P-UR colonies (Figure 5, Table S2).

SBV titers showed behavioral differences in 2 out of 4 comparisons. In contrast with DWV, higher levels in foragers were only observed in older workers from P-UR colonies (P=0.0035) and younger nurses had higher titers in P-R colonies (P=0.035). Interestingly, pollen ingestion was associated with higher SBV titers in younger foragers (P=0.011). On the other hand, age comparisons showed higher titers in older foragers from P-R colonies (P=0.034) (Figure 5).

Behavioral analyses of BQCV only showed significant effects in older workers from P-R colonies, where foragers had higher titers than nurses (P=0.033). In contrast with the other two viruses, younger nurses from P-R colonies had higher titers compared with foragers (P=0.0003). Lastly, in contrast with DWV and SBV, no significant differences were observed in age comparisons (Figure 5).

### 4. Correlations among physiological genes associated with behavior, immune genes and viruses

#### Correlations within categories of genes and viruses

Analyses of the expression of nutritionally-regulated genes associated with behavior revealed strong positive correlations in the age and nutritional comparisons between genes associated with nursing (*vg*-*mrjp1* 100% of age and nutritional comparisons), as well as among genes which expression is associated with foraging (*mfe*-*ilp1* (75%), *mfe-ilp2* (75%), *ilp1-ilp2* (100%). On the other hand, the strongest negative correlation among nursing and foraging associated genes was observed between *mrjp1* and *mfe* (75% of the comparisons) (Figure 6, Table S3).

**Figure 6.**
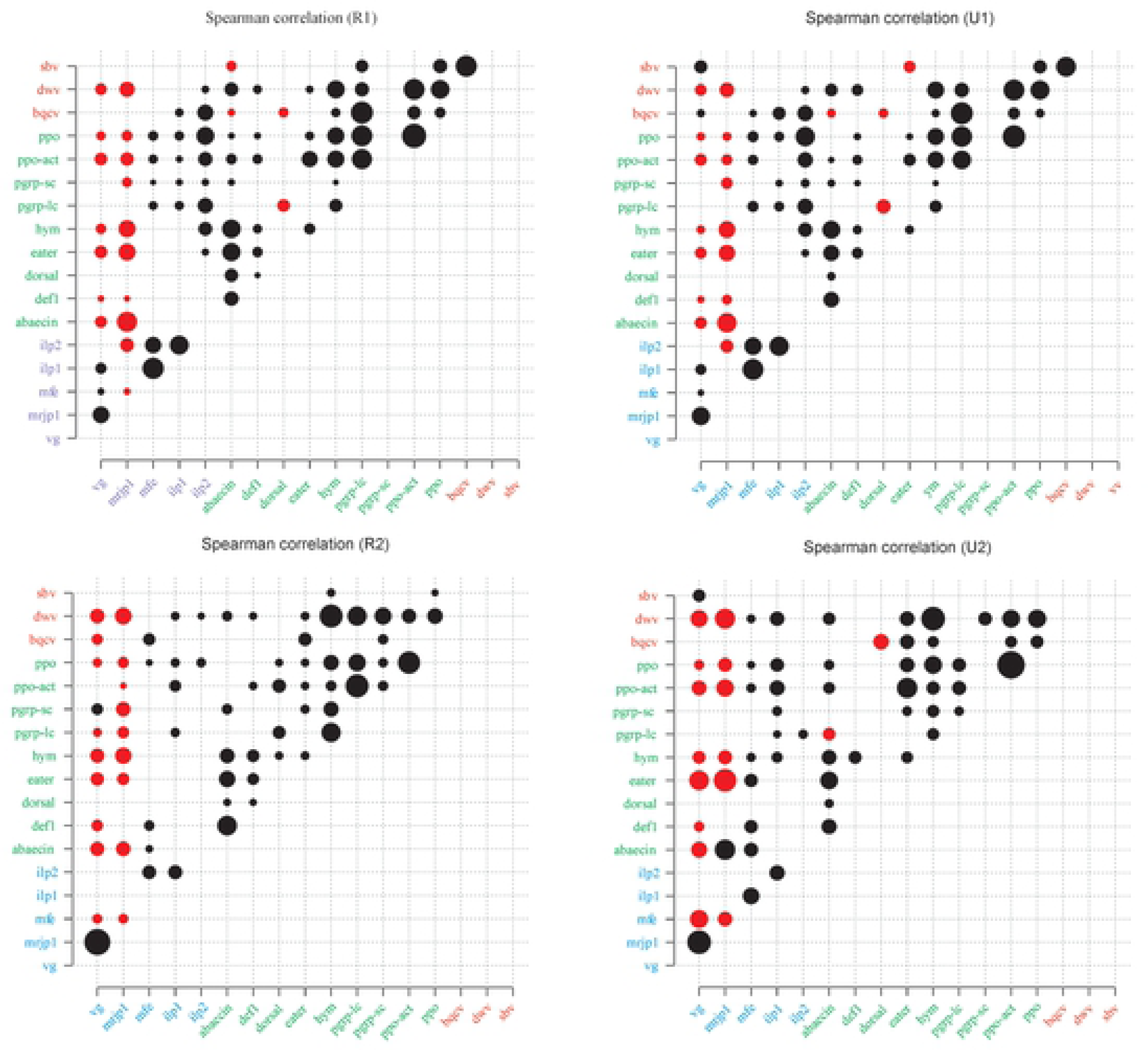
Correlations among physiological genes associated with behavior, immune genes and viruses. Bubble plots representing the correlations between the different variables (genes and viruses) analyzed by the Spearman Rank Correlation Test comparing age and nutritional groups: 2-week-old workers P-R (R-1), 2-week-old workers P-U (U-1), 3-week-old workers P-R (R-2), 3-week-old workers P-U (R-3). Bubbles represent significant associations between the variables (p≤0.05). The size of the bubble represents the R-value and the color represents positive (black) and negative (red) correlations.

As expected, we observed a good association among the expression of the different immune genes analyzed with 61.11 % of positive correlations and 0.7% of negative correlations among gene/nutrition/age comparisons. Among these interactions, we found strong correlations among genes encoding for antimicrobial effectors belonging to the Toll pathway involved in the humoral immune response (*abaecin*, *def1* and *hym*), as well as among genes involved in different functions of the cellular immune response (*ppo, ppo-act*, *eater* and *pgrp-lc*), with 100% of positive correlations among the genes within each group. In contrast, the regulatory gene *dorsal* exhibited the lowest association with other immune genes (28.13% positive correlations). On the other hand, the only significant association among the viruses analyzed was observed between BQCV and SBV, which exhibited a positive correlation in 2-week-old workers (Figure 6, Table S3).

#### Correlations among physiological and immune genes

We first analyzed the relationship among the physiological genes involved in the nutritional regulation of behavior, which expression is associated with nursing (*vg* and *mrjp1*) or forager transition (*mfe*, *ilp1, ilp2*) with immune genes. We find a clear difference in the interactions of immune genes with respect to nursing or foraging associated genes. Thus, the physiological genes associated with nursing showed a strong trend toward negative correlations with immune genes; with *vg* having 66.66% and 2.77% and mrjp1 72.22% and 0% significant negative and positive correlations, respectively, among nutritional and age comparisons. The group of immune genes showing the strongest negative association with nurse associated genes were the three genes encoding antimicrobial effectors (*abaecin*, *def1* and *hym*), followed by the genes involved in the melanization pathway (*ppo* and *ppo-act*), with 100% and 93.75% of negative correlations. On the other hand, the physiological genes associated with the behavioral transition of foragers displayed the opposite pattern, with an average of 41.66% of significant positive correlations, and 0% of negative correlations (*mfe* 44.44 %, *ilp1* 41.66% and *ilp2 38.88 %)*. The genes involved in the melanization pathway showed the strongest positive correlation with genes associated with foraging (79.16%). It is interesting to note that *dorsal* was the only immune gene not showing positive or negative correlations with any of the genes associated with behavior (Figure 6, Table S3).

#### Correlations among physiological genes and viruses

Next, we investigated the relationship between the genes involved in the nutritional regulation of behavior and viral loads. Overall, there is 3X higher percentage of negative correlations (37.5%), between genes associated with nursing and virus levels, compared with positive correlations (12.5%). This trend is mainly a consequence of the negative correlations between *vg* and *mrjp1* with DWV in all age and nutritional comparisons. Although there were positive correlations between *vg* and SBV in workers from P-UR colonies in both age groups, the statistical significance of these correlations is considerably lower compared with the interactions among DWV with *vg* and *mrjp1*. In contrast, the genes associated with foraging showed in general a positive association with virus levels, with 33.33% of positive correlations and 0% of negative correlations. In particular, *ilp2* showed a higher percentage of positive correlations with virus (42.66%), followed by *ilp1* (33.33%) and *mfe* (25%). Virus-specific analyses reveal that both DWV and BQCV exhibit 50% of positive correlations, while SBV does not show significant interactions. The higher percentage of correlations between viruses and physiological genes associated with foraging was observed in the interactions of *ilp2* with DWV (75%) (Table S3).

#### Correlations among immune genes and viruses

Immune genes showed a clear trend toward a positive association with virus infections, with an average of 46.29% of significant positive correlations and 2.7% negative correlations in 108 nutritional and age comparisons. Immune-gene specific analyses revealed the following percentage of positive correlations with viruses: *ppo* (83.3%), *hym* (75%), *pgrp-lc* (66.7%), *ppo-act* (58.3%), eater (41.7%), *abaecin* (33.3%), *def* (33.3%) and dorsal (0%). Virus-specific analysis showed the following percentages of positive correlations with immune genes: DWV (77.8%), BQCV (47.2%) and SBV (13.9%). These results show remarkably specific virus-immune gene interactions, such as those observed among DWV and the genes belonging to the Toll pathway (*abaecin*, *def*, *hym*) as well as with genes involved in the melanization pathway (*ppo*, *ppo-act*), which showed significant positive correlations in all the nutritional and age comparisons. In addition, BQCV showed 100% of positive correlations with *hym* and *pgrp-lc*. Finally, SBV showed little or no correlation with most of the immune genes (0%-25%) with the exception of *ppo* with 75% of positive correlations among nutritional and age groups (Figure 6, Table S3).

### 5. Non-metric multidimensional scaling (NMDS)

An NMDS analysis was performed to visualize the spatial arrangement of the samples according to all the variables analyzed. The best grouping of clusters was two and were nurse and forager bees separately, with no clear differentiation by nutrition or age. Accordingly, behavior was the most correlated variable by itself with the spatial ordination of the samples (p=0.001, R=0.41) followed by its interaction with nutrition (p=0.001, R=0.51) and age (p=0.001, R=0.49). Nutrition was also associated but with lower significance (p=0.04, R=0.06) and age did not show association (p=0.11, R=0.05) (Figure 7).

**Figure 7.**
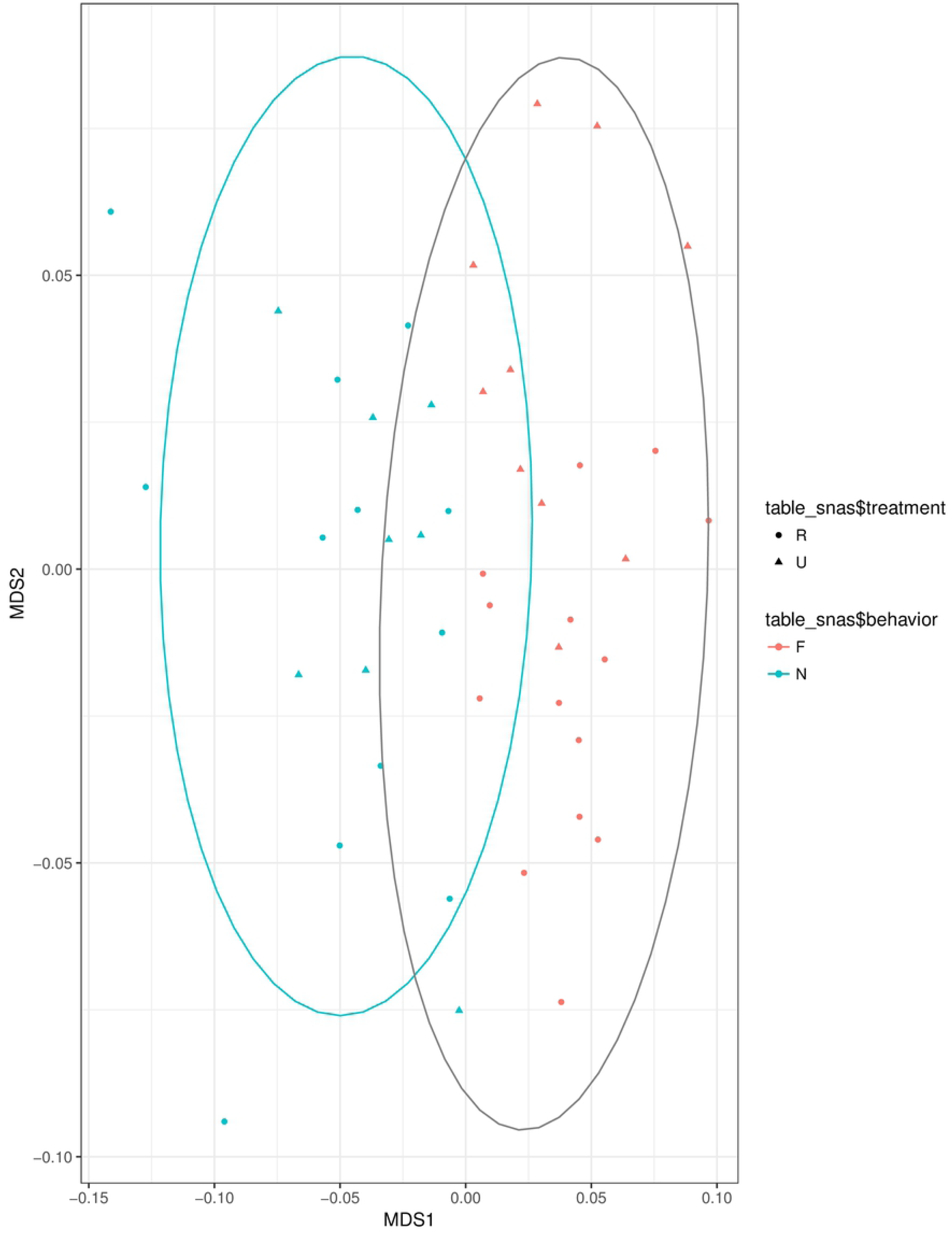
Non-metric multidimensional scaling (NMDS). Visualization of the spatial arrangement of the samples according to nutritional treatment and behavior: P-R (R), P-U (U); foragers (F), nurses (N).

## Discussion

In this study, we used pollen-restricted colonies as a model to study the nutritional conditions experienced in colonies within intensively cultivated agricultural areas. We complemented this experimental model, by using triple TCSC for colony establishment, which allowed us to uncouple the effects of nutrition, behavior and age in colonies of similar size, demography and genetic background. This experimental setup also allowed us to explore the impact of these variables on the expression of physiological genes involved in the nutritionally regulated division of labor, immune genes and three major honey bee viruses. We showed that pollen deprivation leads to a higher rate of new foragers, decreased brood production and caused important changes in the expression of genes associated with physiological activity, immunity and titers of honey bee viruses.

### 1. Effect of pollen restriction on foraging

Our results, showing a higher number of new foragers from colonies with reduced access to pollen, are generally consistent with early studies showing that nutritional stress induces precocious foraging. However, in contrast with other reports, which were conducted under total pollen [48] or honey [26] deprivation, in this study, we investigated the effects of pollen restriction, a more common stress facing honey bee colonies. Other studies where pollen reserves have been experimentally manipulated also had found a negative correlation between pollen reserves and foraging effort [69, 70]. In our experimental colonies, foraging started in both pollen-restricted and pollen unrestricted colonies at the age of 11 days. However, pollen-restricted colonies had a higher number of new foragers compared with non-restricted colonies during the 10-days observation period. Thus, although nurses from both nutritional conditions started foraging at the same day, the higher number of new foragers in pollen-restricted colonies during the full extension of the experiment indicates that the nurses from this group transitioned to foragers at an early age.

By the time the first focal foragers in our study were observed (11 days of age), 25 days after the introduction of mated queens in the hives, newly-emerged bees had already begun to emerge in the colonies. Therefore, it could be expected that the colonies composed by the three cohorts (25, 18, 11-days-old) had a well-established social organization and the behavioral development of the focal bees has not been influenced by demographical abnormalities such as the absence of older individuals (foragers) or young bees. However, in our experimental model, the onset of foraging (at 11 days of age) occurred earlier when compared with typical large colonies, which is around 18.5 days of age [24]. Nurse to forager transition depends on several variables, including population size, nutritional conditions and colony demography [21]. Thus, for example, in small (1000-1200 bees) single cohort colonies established with 1-day old bees and subject to nutritional stress, foraging can start as early as at 3-4 days of age [26]. The onset of foraging observed in our experimental colonies can be due to the interactions of several variables, including small colony size.

We determined the effects of pollen restriction on both nectar and pollen foragers. Our results show a higher number of both types of foragers in pollen-restricted colonies (low pollen storage) compared with unrestricted access to pollen (high pollen storage). Interestingly, although pollen restriction induced new nectar and pollen foragers equally, there was a significantly lower number of new pollen foragers compared with nectar foragers in colonies having unrestricted access to pollen. This result is consistent with early studies showing that stored pollen inhibits pollen foraging [69, 70]. This inhibitory effect of stored pollen along with the fact that brood stimulates pollen foraging [71] have been proposed as mechanisms by which colonies regulate their levels of stored pollen [72]. However, our results, together with other studies [69] showing that pollen restriction resulted in less brood production (see below) and an increasing number of pollen foragers, suggest that the positive association between the amount of brood and foraging requires a determined threshold level of stored pollen. Apparently, below this level, a reduced amount of stored pollen is the main factor stimulating pollen foraging. Altogether, our results indicate that nutritional conditions, associated with pollen storage, have an important effect both on the induction of new foragers and the probability of collecting either pollen or nectar.

### 2. Effect of pollen restriction on brood production and colony demography

In this study, we also showed that there were significantly fewer emerging bees in pollen-restricted colonies when compared with pollen-unrestricted colonies. This result is consistent with previous studies showing a correlation between pollen supply and brood area [69, 73]. It has been proposed that precocious foraging has negative consequences for the demographic balance of colonies, as nurses transitioning early to foragers have a reduced lifespan compared with those that have an extended nurse stage [74–76]. Subsequent forager depletion is expected to result in the further behavioral transition of the remaining nurse population, leading to a reduced capacity to sustain brood rearing, that may be intensified by larval cannibalism [74]. Thus, colonies experiencing nutritional stress due to restricted pollen intake are expected to collapse due primarily to the consequences of worker’s accelerated behavioral development (precocious foraging) on worker lifespan and brood rearing. Our results, showing precocious foraging and reduced brood production in pollen restricted colonies, suggest that a similar situation may occur in regions where colonies surrounded by intensively cultivated areas store low amounts of pollen [14]

### 3. Nutritionally regulated genes associated with the physiological behavioral state

We analyzed the expression of five nutritionally regulated genes associated with the physiological behavioral state. This group of genes include two genes (*vg* and *mrjp1*) associated with nursing physiological stage independently of nutritional condition and age, and three genes (*mfe*, *ilp1* and *ilp2*) whose expression is associated with foraging physiological state when workers are not old or nutritionally stressed.

Our results showed that *vg* RNA levels are affected by behavior, being higher in nurses in all ages and nutritional comparisons. Although high mRNA *vg* levels in nurses are well-known [29, 77], this study provides new information on the effect of age and nutritional conditions on the expression of this gene associated with behavior. Nutritional comparisons between pollen-restricted and pollen-unrestricted colonies showed that increased pollen intake is associated with higher levels of *vg*. Similar results, showing higher *vg* levels associated with pollen feeding, have been obtained in several cage studies [30, 66, 78, 79] as well as in a field colony-level study [14]. However, our study is the first to investigate the effect of pollen feeding in nurse and foragers at different ages at the colony level. Our finding that nutrition influences *vg* expression in most nutritional and age groups, with the exception of older foragers, has practical implications for the use of *vg* as a nutritional biomarker in colony-level field studies. Our expression analysis revealed a trend toward higher expression levels in older nurses irrespectively of nutritional conditions. Although this result was unexpected, a recent study comparing young and old nurses and foragers obtained similar results both measuring mRNA and proteins levels [80]. The biological meaning of this phenomenon it is unknown, however, it is interesting to note that higher levels of Vg are also found in long-lived winter bees [81]. All together, these results reveal a positive association between Vg and longevity, which is compatible with its antioxidant properties [29, 82].

*Mrjp1* expression patterns revealed several interesting insights. First, we found highly significant differences between nurses and foragers independently of nutritional condition and age. Nurse/forager expression differences were especially large in younger 2-week old workers. This expression pattern unveils *mrjp1* as a reliable biomarker of behavior for whole body analyses. Second, young nurses from pollen-restricted colonies had lower *mrjp1* levels compared with pollen-unrestricted colonies (P=0.02), but no significant differences between nutritional groups were found in older nurses. Thus, in contrast with *vg*, the effect of pollen restriction was limited to younger nurses. Third, foragers showed intriguing changes of *mrjp1* expression associated with nutrition and age. Younger 2-week-old foragers from colonies with high pollen storages (P-UR), have higher *mrjp1* levels compared with foragers from colonies with low pollen storage (P-R). On the other hand, older foragers (3-week-old) experienced an increase in both nutritional groups compared with younger foragers (2-week-old).

What can be the biological meaning of these results? *Mrjp1* is expressed in different tissues and organs besides the hypopharyngeal glands [64, 83, 84], suggesting that in addition to their traditional role as a secreted protein, it is used endogenously for other nutritionally related functions [83–85]. An initial step toward the clarification of the biological meaning of these results it is to determine whether these high levels of *mrjp1* expression are the consequence of increased expression in the hypopharyngeal glands or other organs with non-exocrine function. Still, these results suggest interesting possibilities for future studies. Royal jelly proteins secretion from nurses’ hypopharyngeal glands has been primarily associated with larval and queen feeding. However, these proteins also are transferred during trophallaxis – mouth to mouth food transfer – preferentially to younger workers [86, 87]. Consistently, our results suggest that foragers could be MRJPs donors during trophallactic interactions and that the potential for these interactions increases with nutrition and age. Interestingly, it has been hypothesized that the transfer of royal jelly proteins among workers could provide information about colony pollen reserves and act as cue for regulation of foraging [88].

Our results provide interesting insights into the dynamics of *mfe* expression in whole-body analysis. First, we found higher levels of *mfe* in foragers compared with nurses in younger workers of colonies not experiencing nutritional stress. However, it is interesting to note that the magnitude of this difference is lower in contrast to results obtained by direct comparison of isolated corpora allata from nurses and foragers [89]. This result suggests that *mfe* expression in other tissues of young nurses reduces the difference in overall body expression. Consist with this possibility, *mfe* expression has been detected in ovaries of nurses honey bees [84] and JH synthesis have been reported in ovaries of beetles [90] and mosquitoes [91]. An additional report showing higher levels of mfe in abdomens of young nurses compared with young foragers from a non-nutritionally stressed colony, is further consistent with the proposal of that *mfe* is significantly expressed in other tissues different to the corpora allata, especially in young non-nutritionally stressed nurses [80], although still forager’s corpora allata is the main organ for JH synthesis. Thus, these results also contribute to challenge the traditional view that the corpora allata is the only source of JH in insects [33, 92, 93]. On the other hand, our nutritional comparison showed a trend associating higher *mfe* levels with pollen consumption, but significant differences were only observed in older workers. Lastly, the expression of this gene significantly declined in older foragers from pollen-restricted colonies. Overall our results showed behavioral differences in the expression of *mfe*, but also revealed that expression is inhibited in various degrees by nutritional stress and age.

Previous studies have shown that insulin-like peptides (*ilp1* and *ilp2*) are involved in the nutritionally mediated regulation of division of labor in honey bees [27, 30]. Insulin-like peptides are expressed in different tissues and organs including the brain [29, 30], fat bodies [27] and ovaries [84]. Although these studies have revealed the existence of differential tissue-specific responses to nutritional inputs, few tissues and organs have been so far analyzed and it is unknown the contribution of their expression relative to the whole body. Our analysis of *ilp1* showed that this gene presented behavioral differences in 2-week-old workers from colonies with unrestricted access to pollen, in which foragers had significantly higher levels compared with nurses. Thus, *ilp1* expression patterns in the whole-body followed a similar pattern to that reported in isolated brains, but this similarly is restricted to young individuals of colonies not experiencing nutritional stress. The magnitude of the difference between nurses and foragers for the whole body is lower than that observed in brains [29, 30]. This result suggests that in young nurses *ilp1* expression is upregulated in other tissues when the colony is not nutritionally stressed. This explanation is consistent with the report of up-regulation of this gene by amino acid supplementation in fat bodies of 7-days-old bees [27] and with the observation of higher level of *ilp1* in the abdomens of young nurses compared with young foragers in non-nutritionally stressed colonies [80]. On the other hand, given the expression differences yet existing between young nurses and foragers from colonies not experiencing pollen deprivation, our results also suggest that under this nutritional condition, *ilp1* is predominantly higher expressed in forager’s brain compared with other tissues of nurses (e.g., fat body). In addition, nutritional comparison among young foragers indicates that *ilp1* expression is reduced by nutritional stress, possibly by inhibiting *ilp1* forager’s brain expression. In contrast, in older nurses, nutritional stress is associated with increased expression of *ilp1*. It remains to be addressed the contribution of the brain and other tissues in this response. On the other hand, age comparisons revealed an important age-dependent reduction in its expression in both nutritional conditions. Thus, *ilp1* follow similar mRNA levels and expression pattern compared to *mfe*, but with some differences in the strength of the effects of nutrition and age.

The expression profile of *ilp2* showed interesting similarities and differences compared with *ilp1*. In contrast with *ilp1*, *ilp2* showed higher levels in young foragers in both nutritional conditions. Previous brain expression analyses revealed no behavioral differences in *ilp2* expression [30] suggesting that *ilp2* expression in other young foragers’ tissues could account for the observed differences in this whole body analysis. On the other hand, similarly, to *ilp1*, significant differences between nutritional groups were only observed in older workers.

The finding of no expression differences of *ilp2* associated to nutrition in young nurses, is consistent with a study reporting no nutritionally related differences in fat body *ilp2* expression in young nurses at colony level [27]. However, this result contrasts with the report of higher expression levels of *ilp2* associated with pollen ingestion in an abdominal transcriptome analysis [63]. Age comparisons showed even more contrasting differences in the expression of both genes: while *ilp1* showed a general age-dependent down-regulation, *ilp2* was upregulated in workers from pollen-restricted colonies.

### 4. Immune genes

Nutritional treatments had a significant effect on 1/3 of the behavioral and age gene comparisons. However, in most comparisons showing expression changes associated with pollen availability, the expression of the immune genes was upregulated by pollen ingestion compared with pollen restriction. Moreover, it is interesting to note that most of the comparisons not showing up-regulation by pollen ingestion were observed in a single non-effector regulatory gene (*dorsal*). Thus, despite pollen availability having a discrete effect on the expression of immune genes, the results obtained are in general consistent with the notion that availability of nutritional resources is required to keep an active immune system [94]. From another angle, our result also revealed that dorsal was the immune gene most up-regulated by nutritional stress. This finding is significant as this is a regulatory gene that activates the expression of effector immune genes belonging to the Toll pathway [52, 95]. Thus, it remains to be determined the mechanism by which this gene react to nutritional conditions and its possible interactions with the IIS/TOR pathway involved in the nutritional regulation of behavior.

Analyzing same-age nurses and foragers is considered a gold-standard for comparing task-dependent differences [96]. Our experimental design allowed us to examine the expression of immune genes in nurses and foragers of the same age at two different time points, uncoupling the effects of behavior and age on the expression of different types of immune genes for the first time. Changes in immune function had been associated with a behavioral transition in honey bees. Foragers experienced an important reduction in the number of total hemocytes, an effect reproduced by JH treatments [97, 98]. These results lead to two proposals: first, a reduction in the abundance of circulating immune cells in foragers could imply that immunity is a function of behavioral caste [98, 99]. Second, honey bee colonies could economize nutritional resources by downregulating the forager immune system [98–100]. Our results showing higher expression in foragers are consistent with the proposal that the nurse to forager transition is associated with changes in honey bee immune function. However, these results are inconsistent with the view of an immune system downregulation associated with forager behavior. It remains to be determined the mechanisms responsible for foragers’ increased expression of immune genes. It is possible that this higher expression of immune genes is the result of increased contact with pathogens during foraging. Alternatively, the higher expression of immune genes could be the result of a programed genetic response regulated by nutritional clues to prepare the forager for a higher probability of encountering pathogens outside the hive. This proposal implies that the colony does not economize energetic resources by downregulating the immune system in foragers.

Another interesting insight into the expression of immune genes is that age had an important effect on their expression. The regulatory gene *dorsal*, was the immune gene most strongly up-regulated by age, although two genes encoding for antimicrobial peptides (*def1* and *abaecin)* and the immune genes encoding pattern recognition proteins (*pgrp-lc, pgrp-sc*), also showed significant upregulation in at least one behavioral/nutritional group. In contrast, the genes involved in the cellular immune response did not show increased expression with age (*ppo*, *ppo-act*, and *eater*), but rather nutritionally and behavioral dependent changes in their expression, with consistently lower expression in foragers from pollen-restricted colonies, and inconsistent changes in the expression among these genes in nurses and foragers from pollen-unrestricted colonies.

Our results suggest that while the activation of immune response after the nurse to forager behavioral transition involves both the humoral and cellular immune response, age-dependent changes in their expression are partly behavioral and nutritional dependent, with old foragers being unable to maintain high expression of genes involved in the cellular immune response when experiencing nutritional stress. Interestingly, reduced expression of genes involved in the cellular immune response also has been reported in overaged winter bees [101]. On the other hand, Bull et al., (2012) reported increased expression of genes involved in the humoral immune response in old foragers compared with 1-day old bees as well as increased foragers’ resistance to fungal infections. These results lead the authors to infer that reduced pathogen susceptibility in forager bees is associated with age-related activation of specific immune system pathways [102]. While the comparison between young bees and old foragers in that study, did not allow differentiation of the effects of behavior and age, the results are consistent with our findings of higher expression of immune genes in foragers and the up-regulation of regulatory and effector genes involved in the humoral immune response in older bees. Nevertheless, our results do not support the view of a general upregulation of immune gene expression in older bees, especially those related to the cellular immune response. On the other hand, our results neither support the view of a decline in the immune competence associated with the humoral immune response. Interestingly, a recent study analyzed the effects of bacterial injections of the Gram-negative bacterium *Serratia marcescens* on the expression of *def1* both using nurses and foragers (unknown age) from a typical colony, as well as young (6-8 days-old) and old (21-13 days-old) nurses and foragers using a single cohort colony [80]. Consistently with our results of the humoral immune genes, it was found higher expression of this gene in foragers compared with nurses independently of the age, behavioral status and type of colony analyzed. In addition, analyses of the fold-change in the expression levels of the bacterial injected bees relative to the controls, showed stronger immune response in nurses compared with foragers. Although this last result was interpreted as an evidence of immunosenescence, the higher expression levels of foragers compared with nurses, which are even stronger at older age, makes this interpretation controversial.

Previous studies have analyzed behavioral related differences in different aspects of immune function. Two potential confounding factors in some of these studies are the comparisons of foragers of unknown age and newly-emerged workers, which are considered nurses. The comparison of foragers of different age could not take in consideration existing age-related changes in gene expression. On the other hand, newly-emerged bee are not mature nurses and, consequently, the expression of a wide range of genes involved in metabolism and physiology is considerably different than in functional nurses [29, 80, 103]. Despite these potentially confounding factors, our results are in general consistent with previous studies reporting higher PO and AMP activities in foragers compared with young bees [104, 105]. Of particular interest is the study by Schmid et al., (2008), who compared nurses and foragers of different physiological ages (5-11 days-old age-right nurses and precocious foragers, 23-28 days-old age-right foragers and overaged nurses). Despite the finding of higher PO activity in age-right foragers compared with age-right nurses, the number of total hemocytes was significantly lower in age-right foragers. In addition, age comparisons showed an important decline in hemocyte number in older nurses and foragers; although overaged nurses have marginal, but significantly, higher hemocyte numbers compared to age-right foragers of similar age. These results suggest that overall hemocyte decline is mainly age-dependent, but that behavioral state also may have an additional effect on this process. Based on these results, the authors proposed an age-dependent abandonment of cellular immunity in the honey bee [100]. However, this view is apparently missing some elements to explain the important discrepancies observed between total hemocyte number and PPO enzymatic activity. Thus for example, why age-right foragers (low hemocyte number), have higher PPO activity compared with age right nurses (high hemocyte number)?

In contrast with other insects [52, 106, 107], the honey bee genome harbors a single gene encoding PPO [52, 108]; but similarly to other arthropods [107, 109], the honey bee *pro-ppo* lacks a signal peptide for extracellular exporting [52]. Interestingly, this lack of a signal peptide in the *pro-ppo* gene implies that PPO is not released by secretion but by lysis of the hemocyte [61, 107, 109]. This characteristic of PPO’ function could be important to explain some aspects of honey bee cellular immune response. At present, it is not clear if hemocyte lysis-mediated PPO release occurs in continuous or discontinuous mode during adult bee ontogeny and how it is affected by behavioral state, age, nutrition and immune challenges. Still, PPO release after the noticeable reduction in total hemocyte number, could explain why PPO activity remains high in foragers [100, 104, 105] and the observation that immune challenges resulted in reduced PO activity, suggesting a failure in replenishing synthesis of PPO [105] possibly associated with hemocyte lysis. However, increased PO activity in old foragers is also consistent with the proposal that the persistence of active cellular immune function after the reduction of total hemocyte number results from the selective reduction of the of non-immune hemocyte population (prohemocytes) and the conservation of hemocytes with immune activity (granulocytes and plasmatocytes) [104]. Our expression data of the cellular immune genes involved in the melanization pathway, even when showing reduced expression levels in old workers, still revealed high expression in old foragers compared with old nurses as well as changes in their expression associated with nutrition. We propose two alternative explanations for our results regarding the expression of genes involved in the cellular immune response, which still can be compatible with the above-mentioned reports. First, there is a reduction of total hemocytes mainly associated with age [100]. This is consistent with the observed decrease on the expression of cellular immune response genes in old workers [102] as well as in winter bees [101]. This hemocyte reduction mainly occurs in non-immunogenic hemocytes [104], but also there is a modest reduction in immunogenic hemocytes. This remaining population of hemocytes in old foragers experience important changes in their expression of cellular immune genes primarily associated with behavior and secondarily with nutrition. Alternatively, the observed reduction in total hemocytes, may do not involve a significant reduction of immunogenic hemocytes [104], but rather a reduction on their expression of cellular immune genes also associated with age, in addition with behavior and nutrition.

#### Virus levels

A notable result in our study is the finding that behavioral state had an important influence on viral titers. This behavioral effect was stronger in the case of DWV, followed by SBV, while BQCV was the virus less affected by behavior. DWV and SBV have higher viral titers in foragers in most of the nutritional and age groups. Our results show that the only nutritional and age groups not showing behavioral differences on these viruses are the younger workers from non-pollen restricted colonies. This suggests that both nutritional stress and older age contribute to increase viral titers in foragers.

The finding of higher DWV levels associated with foraging has important implications for the understanding of the relationships among nutritional stress, behavioral development and colony losses caused by viral infections. Nutritional stress caused by pollen dearth is a common situation experienced by colonies within intensively cultivated agricultural areas [14, 66, 79]. Pollen deprivation induces precocious foraging, which in turn is associated with accelerate aging and reduced lifespan [75, 76]. We anticipate the possibility that increased DWV titers associated with foraging could further reduce forager survival and exacerbate the changes in colony demography leading to colony collapse [65, 74]. In this study, we were unable to precisely estimate the demographic composition of the experimental colonies during different time collections. However, the number of nurses and foragers that we collected at each time point provides an approximate estimation. It is noteworthy that older 3-week-old foragers from pollen-restricted colonies were 2.72 times more abundant compared with foragers from colonies with full access to pollen. In this regard, it is interesting to note that the highest DWV titers were observed in 2-week old foragers from colonies with full access to pollen. A possible explanation for this result is higher mortality in foragers from colonies with unrestricted access to pollen.

Another interesting result in our study is the finding that pollen ingestion was associated with significantly higher levels of DWV and SBV, observed in younger nurses and foragers, respectively. In fact, BQCV was the only virus showing higher titers associated with pollen restriction, with higher levels in younger nurses. There were interesting virus-specific interactions of nutrition with behavior and age. First, although direct nutritional comparison between young nurses showed higher DWV titers associated with pollen ingestion, significantly higher titers in younger foragers were only observed in pollen-restricted colonies. Second, age shows to influence the effect of nutrition on viral titers, as significant nutritional differences were only observed in relatively younger (2-weel-old) workers.

The effect of nutrition on DWV levels has been previously analyzed in different studies using caged bees in laboratory conditions [63, 65, 67]. Our results revealing an association between pollen ingestion and higher DWV titers at the colony level contrast with an initial report showing that pollen ingestion decreases viral replication in comparison with undernourished bees [65]. However, new lines of evidence are consistent with our results. First, DWV-B replication is inhibited when there is co-infection with *Nosema ceranae* [67] suggesting that competition for nutritional resources reduces viral replication. Second, Alaux et al., (2011) found higher DWV titers associated with pollen feeding in non-varroa parasitized bees. Moreover, these authors also found that pollen feeding further increases viral replication in varroa-parasitized bees already having high DWV levels. This suggests, not only that nutritional resources increase DWV replication, but also that this effect is even stronger in colonies with high varroa infestation levels and concomitant increased viral loads. Based on these conditions, high nutritional status is not only no beneficial, but detrimental for survival [63]. Our results suggest that similar nutritional and viral interactions take place at the colony level. However, we were unable to establish an association between Varroa level and DWV, since we did not measure varroa infestation levels in our experiment. Still, it is reasonable to expect that our colonies experienced moderate infestation levels since no acaricide treatment was applied after the colony’s establishment in early summer.

Comparisons between 2-week and 3-week-old workers showed discrepant effects of age on the titers of the viruses analyzed. While DWV showed significantly lower titers in older bees in workers from colonies with unrestricted access to pollen, SBV showed increased infection levels in foragers from pollen-restricted colonies and no age-related changes were observed in BQCV virus. DWV was the virus most affected by age, with a trend to lower titers in all the comparisons, although significant lower titers in older workers were only observed in colonies with full access to pollen. The observed lower DWV titers in older foragers have two main possible explanations. First, an antiviral immune response could be triggered resulting in lower titers in older bees. Second, alternatively, our result could show a preferential survival of workers with lower DWV titers.

#### Correlations

Our analysis of significant correlations among the expression of the different group of genes and viruses analyzed revealed clear trends in their interactions. As expected, analysis of the targeted gene categories revealed strong positive correlations among the physiological genes associated with nursing as well among the genes associated with foraging, confirming the co-regulation of both groups of genes and supporting their use as molecular markers of behavior. Similarly, a general correlation was observed among the expression of the immune genes, with particularly strong positive association among the genes belonging to specific immune pathways, such as the genes encoding AMP and genes involved in the melanization pathway. These results confirm the coordinated response of the different immune pathways, which are co-regulated in a similar manner primarily by behavioral-associated physiological changes, but also by nutrition and age. In contrast, the titers of the three viruses analyzed did not exhibit significant correlations among the different nutritional, behavioral and age comparisons, suggesting that these factors differentially influence their replication.

On the other hand, the groups of physiological genes associated with behavior showed a clear difference in their interactions with immune genes. Thus, while the genes associated with nursing showed a strong negative correlation with immune genes, the genes associated with foraging displayed the opposite pattern (Figures 6 and 7). An interesting exception was the regulatory immune gene *dorsal*, which *did not* show correlations with the genes associated with behavior. These results further support the view of an association between the physiological pathways involved in the regulation of division of labor with the immune system. Moreover, these results confirm the upregulation of effector immune genes in foragers, independently of nutritional conditions or age.

The physiological genes associated with behavior also showed differences in their interaction with DWV and BQCV. In particular, nursing associated genes showed a strong negative correlation with DWV, while foraging associated genes showed a predominant positive correlation (Figure 6). A notable exception among these interactions was observed in SBV, which did not show significant correlations with behavioral-associated genes.

Finally, our analysis of the interactions among immune genes and honey bee viruses revealed a predominant positive association between the expression of most of the immune genes and virus infections. In particular, DWV showed a strong positive correlation with genes encoding for AMPs and those involved in the melanization pathway (100%). In contrast, SBV exhibited a low percentage of positive correlations with most immune genes (0-25%), with the exception of *ppo (75%)*. Similarly, *dorsal* was the immune gene which expression showed the lowest number of correlations with viral infection. Although these results revealed a strong association between the infections of two of the most common and prevalent honey bee viruses in European honey bees, DWV and BQCV, and the expression of immune genes, our analysis is unable to distinguish the mechanisms by which these variables are associated. Thus, it is either possible that the higher expression of the immune genes observed in foragers can be a response to increased DWV infection, or that this immune expression occurs before the increase in viral infection. Still, the close association observed between *ppo* expression and DWV titers, which has been documented in winter bees (Steinman 2015) is remarkable and consistent with the association between the cellular immune response and viral infections reported previously in honey bees and other insects [110–113].

#### Concluding Remarks

In this study, we used pollen-restricted colonies as a model to study the nutritional conditions occurring in colonies surrounded by intensively cultivated agricultural areas which typically experience nutritional stress caused by pollen dearth. This experimental model was complemented by the use of a new technique for colony establishment that allowed us to uncouple the effects of behavior, age and nutrition. Our results revealed that pollen restriction induced early behavioral transition to foraging and that this behavioral change to foragers is associated with a general increase in the expression of immune genes. The expression of immune genes also shows a trend towards increased expression associated with pollen ingestion and an age-dependent regulation of specific immune pathways, including a sustained high expression of genes involved in the humoral response and down-regulation of genes involved in the cellular response in old workers. An interesting exception was *dorsal*. The expression of this regulatory gene was the less affected by behavior and the most affected by nutrition. Although, dorsal expression still it was up-regulated in foragers, in contrast with other immune genes, its expression was mostly upregulated by nutritional stress. Since dorsal is a regulatory gene involved in the regulation of the expression of effector immune genes, its observed sensibility to bee’s nutritional conditions suggests that this gene has an key integrative function on the interactions among nutrition, nutritionally-dependent behavioral development and immune response.

The analyses of three predominant honey bee viruses (DWV, BQCV and SBV) show that they are affected to various degrees by honey bee behavior, nutrition and age. Our main results include the demonstration of higher DWV levels in foragers and unexpectedly, the finding that pollen ingestion is associated with higher DWV in young nurses. Higher DWV in foragers suggests that accelerated behavioral development induced by nutritional stress could lead to amplified forager mortality and further exacerbate changes in colony demography associated with colony collapse. On the other hand, high DWV levels associated with pollen ingestion, suggest that in infected colonies, nutritional resources further enhance the replication of this virus.

Our analyses of the correlations among the expression of immune genes and viral titers with physiological genes associated with behavioral state revealed that the immune genes and DWV exhibited strong negative correlations with genes associated with nursing, but positive correlations with genes associated with foraging (Figure 8). These results emphasize the role of nutrition on worker behavioral development and their consequences for honey bee immunity and susceptibility to pathogenic infections. Finally, the results obtained by the NMDS analysis further support the conclusion that behavioral state is the most important factor associated with the expression of immune genes and viral titers. Overall, we provide important insights into the mechanisms by which nutrition, behavior and age alter the expression of key genes involved in the regulation of honey bee physiology and immunity, which are expected to contribute to a better understanding of the effects of nutritional stress on colony health.

**Figure 8.**
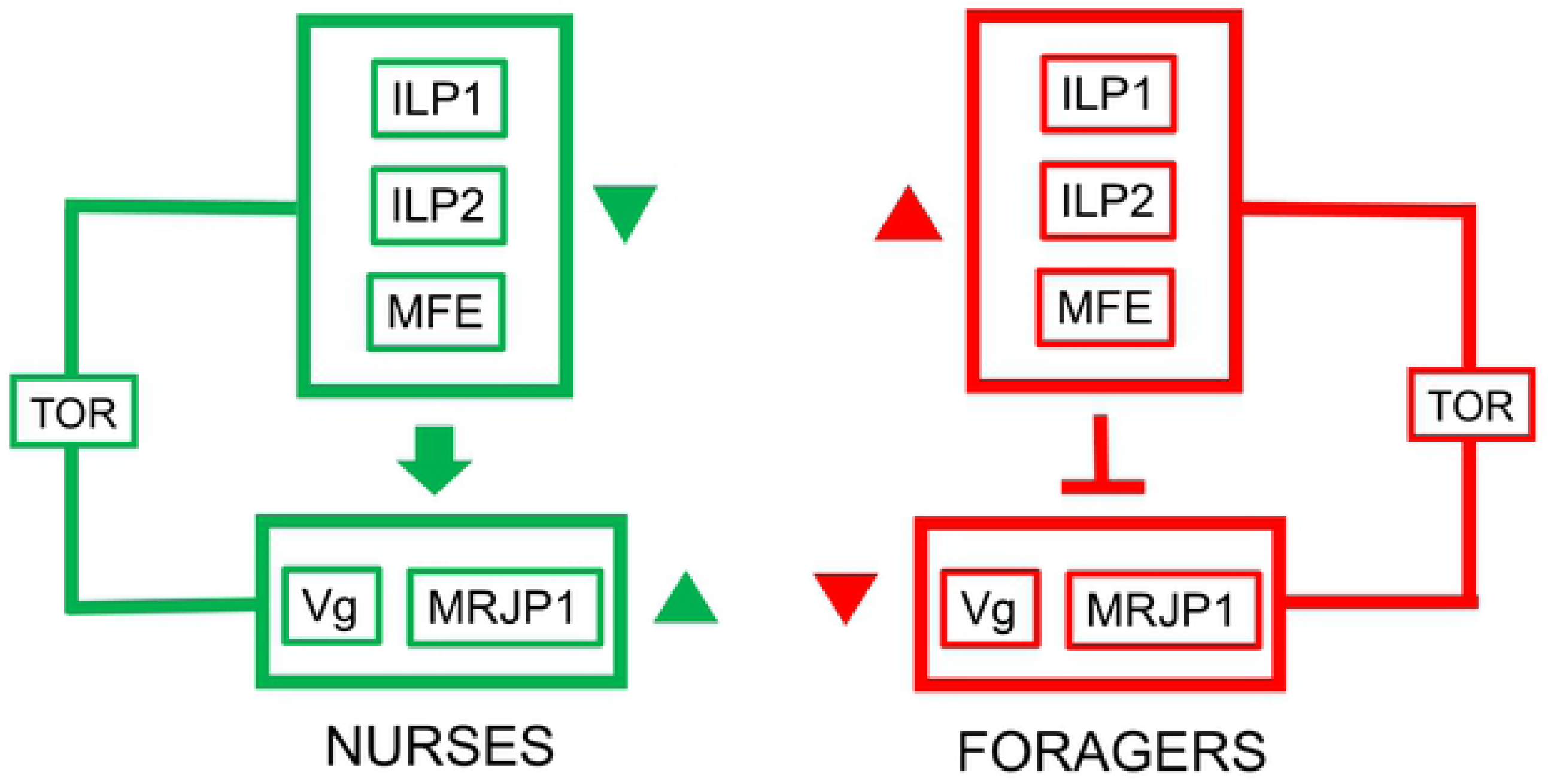
Schematic representation of the predominant correlations of whole-body levels of nutritionally regulated physiological genes, immune genes and DWV associated with behavioral state. Up-regulated and down-regulated levels between nurses and foragers are represented by upward and downward pointing arrowhead, respectively.

## Materials and methods

### 1. Experimental Colonies

Field experiments were performed in the apiaries of the USDA-ARS Bee Research Laboratory in Beltsville Maryland. The triple cohort composite colony (TCC) is a technique of colony establishment originally designed to vary colony age demography in colonies of similar size and genotypic structure. We created triple cohort sequentially-composed colonies (TCSC) by modifying a previously described protocol [114]. Instead of simultaneously introducing three cohorts (nurse and foragers of unknown age, plus newly-emerged bees), we introduced sequentially (weekly) three groups of newly emerged bees during a period of two weeks. This modification resulted in colonies with a population of three thousand bees composed predominantly of 2-week-old foragers, 1-week-old nurses, and newly-emerged bees allowing us to uncouple the effects of age and behavior by collecting same-age nurses and foragers at different times. Then, in contrast with the original protocol with colonies headed by non-egg-laying queens confined in small cages, we confined egg-laying queens in frame-sized excluder cages. This last modification avoided the necessity of introducing young bees into the colony to prevent delayed behavioral development [114] (Figure S1).

A set of sixteen TCSC colonies, each with a population of three thousand bees were formed by newly-emerged bees from multiple colonies (approximately 50) and established in four-frame boxes. This experimental design homogenizes the genetic background of the colonies studied: although the genetic variation within colonies is high, variation across the different colonies is expected to be low. For each cohort, newly emerged bees were collected from combs of mature pupae taken from colonies derived from populations of *Apis mellifera ligustica* and placed in an incubator at 32°C and 55% RH. Newly emerged bees were thoroughly mixed, marked on the thorax with enamel paint, and distributed into the colonies. After establishing the colonies, two experimental nutritional groups were formed: pollen restricted (P-R) and unrestricted (P-UR) colonies (N=8 per group). P-R colonies were formed by removing the frames containing pollen and by activating the pollen traps. In P-UR-colonies the frames with pollen were not removed and the pollen traps were not activated. We performed subsequent behavioral and molecular analysis in the last cohort of newly-emerged bees.

### 2. Behavioral Observations

Daily monitoring to detect foraging was initiated five days after the introduction of the last cohort. This time is considered the minimum required for nurse to forager transition [26, 74]. Behavioral observations began when the first foragers were discovered and continued for ten consecutive days. Each colony was observed for three periods a day (10 AM-12 PM, 12-2 PM, 2-4 PM). In each period, the hive entrance was blocked with a metal screen for 10 minutes, followed by a 5-minute observation time. Returning pollen and nectar foragers were identified by the observation of pollen loads on their curbiculae or the distension of their abdomens, respectively [26]. The average quantity of pollen captured in the pollen traps in the P-R colonies, after the establishment of the colonies and until the end the observations, was 40.4 grams (28.93 to 44.93 grams).

### 3. Quantification of the number of emerging bees and collected pollen

Two days after the end of the behavioral observations, and one day after the last collection of 3-week-old workers for molecular analysis, pollen trapped on P-R colonies was collected and the frames containing brood (one frame per colony) were removed from all the hives, transferred to the laboratory and maintained at 32°C in an incubator. Emerging bees from P-R and P-UR colonies were counted until the last bee emerged after 16 days.

### 4. Sample collections for molecular analysis

Collections were performed two (2W) and three weeks (3W) after the last cohort was introduced into the colonies. In each of these two age collections, nurses (N) and foragers (F) of the same age were collected from P-R and P-UR colonies. Therefore, eight age, behavioral and nutritional groups were collected for subsequent analysis. The total numbers of bees collected per each of these groups were as follow. 2W: P-R N (n=76), P-R F (n=78), P-UR N (n=76), P-UR F (n=53). 3W: P-R N (n=53), P-R F (n=53), P-UR N (n=44), P-UR F (N=23). Foragers were collected from the entrance of each colony (double painted) and nurses from the brood area (single painted). Immediately after their collection, bees were flash-frozen in liquid nitrogen for subsequent RNA-extraction and gene expression analyses.

### 5. Gene expression analysis

Whole body bees were homogenized in Trizol (Invitrogen) in a Fast-prep-24 (MP) instrument. Whole bodies were used for analysis because different tissues and organs are expected to participate in the regulation of behavior, immune function and susceptibility to pathogens. RNA extractions were performed with the RNeasy extraction kit and genomic DNA traces were removed using DNAse I (Qiagen). RNA was eluted in 100 μL. For each sample, cDNA was synthesized using 1 μg of total RNA and Thermo-fisher reagents, including M-MLV Reverse Transcriptase (40U), RNAse inhibitor (25U), random hexamers (2.5 uM) and dNTPs (0.8 mM) in a final reaction volume of 25 μL. Thermal profile for cDNA synthesis was as follows: 25°C (10 min), 48°C (45 min) and 70°C (5 min). Each cDNA reaction was diluted by adding 100 μL buffer (10 mM Tris HCl pH 8.5. Transcription levels were quantified by qRT-PCR using a Life Technologies Vii7 system, with SYBR green reagents and a two-step thermal profile for amplification (95°C 15 sec, 60C 1 min). All assays were performed in triplicates in a final volume of 10 μL. Data from multiple plates were normalized with a reference sample and *Rps5* as an internal control gene [115] using the *Gene Study* software package (Life Technologies). Quantification of mRNA levels was performed by the ΔΔCT method [116]. Relative values were calculated based on the differences (ΔCT) between the CT values of the focal gene/virus and *Rps5* and then transformed to relative values. For each comparison, ΔCT values exceeding 2 SD from the mean were considered technical outliers and removed from the analysis. Primers were designed using the Oligo 7 programs [117] and their efficiencies tested using a relative standard curve from concentrated cDNA (Table S1).

### 6. Statistical analyses

Initial analyses of normality and homoscedasticity of the variables was carry out using the Kolmogorov-Smirnov and Levene tests [118–120]. Differences between groups of the variables that fulfilled the assumptions of parametric statistics were analyzed with ANOVA and Scheffeé tests and non-parametric variables were analyzed by Kruskal Wallis and Mann Whitney U test [121]. On the other hand, correlations in the expression level of the different genes were analyzed considering nutritional groups by the Spearman Rank Correlation Test [120, 121]. Spatial representation of all samples and variables was visualized with a non-metric multidimensional scaling (NMDS) and the best number of clusters from this representation was determined with the function NbClust using the Euclidean distance and k-means method. The association between nutrition, behavior, age and the interaction between them with the ordination of the samples was determined with the envfit function [120, 122]. In all cases, p-values under 0.05 were considered statistically significant.

## Acknowledgments

We are grateful to Bart Smith for his advice and important collaboration in the establishment of the experimental colonies. The authors would like to thank Madlin Rizkalla, Alicia Santiago, Danielle Baker and Dawn Lopez for assistance in the collection and processing of the samples. We also acknowledge Vanessa Corby-Harris, Rosalind James and Eugene Ryabov for reviewing the manuscript.

## Author Contributions

Conceived and designed the experiments: MC. Performed the experiments: MC SM. Analyzed the data: MC BB SM JE YC. Contributed reagents/materials/analysis tools: MC JE YC. Wrote the paper: MC BB SM JE.

## Funding

This work was supported by the ARS-USDA funds granted to the Bee Research Laboratory.

The authors declare non-competing interests

## Supplementary Figures

**Figure S1. The triple cohort sequentially composed colony (TCSC) model for colony establishment.** All colonies were founded within a 2-week period in three consecutive steps. The first cohort of one thousand newly emerged bees was introduced into the hives together with the queens. Then, subsequent batches of one thousand newly emerging bees were introduced after one week (second cohort) and two weeks (third cohort). At the beginning of the colony establishment, each hive was provided with one frame of honey and one frame of pollen. After establishing the colony, we performed subsequent behavioral and molecular analysis in the last cohort of newly emerged bees. This final cohort was expected to experience normal behavioral development, given the previous introduction of two older cohorts into the colony and the subsequent incorporation of newly emerged bees. The behavioral development of the final cohort was monitored to determine the onset of foraging and then sample collections were performed at the age of 2 and 3weeks.

**Figure S2. Bihourly number of new foragers.** Upper panel (A) pollen foragers. Lower panel (B) nectar foragers. The y-axes represent the daily mean of both types of foragers during different bihourly observations: times are indicated using the 24-hours daily system. 10-12 hours, 12-14 hours, 14-16 hours. The x-axes indicate the age of the new foragers. Restricted colonies (orange line), unrestricted colonies (blue line). Error bars represent SE across colonies. Daily comparisons of the number of new nectar and pollen foragers during the three time periods of the observations (10-12, 12-14 and 14-16 hrs.) showed higher time-related variation in the number of nectar foragers compared with pollen foragers. Nectar foragers exhibited higher differences between P-R and P-UR colonies during the first observation period (10-12 AM) compared with the last observation (14-16 hrs.). In contrast, pollen foragers showed variation among bihourly observations only during the first three days. Afterward, they followed a similar pattern in the three observations periods, characterized by a significantly higher rate of foraging in P-R colonies

**Figure S3. Daily number of new foragers.** Daily mean of new foragers from both nutritional conditions (y-axis) according to their age (x-axis). Error bars represent SE across colonies. Pooled daily analysis of the three observation times exhibited a significant number of new foragers in P-R colonies starting from day 13 until the end of the observations on day 20.

## Supplementary Tables

**Table S1. Primer pairs used for quantitative PCR of honey bee genes and viruses.** Primer’s directions are included at the end of their respective names: Forward primer (F). Reverse primer (R).

**Table S2. Statistical results of nutritional, behavioral and age comparisons of individual genes and viruses.** Different font colors represent relevant behavioral (red), nutritional (green) and age (blue) comparisons.

**Table S3. Statistic results of correlations among honey bee genes and viruses’ levels**. P and R-values are classified as following: vertically, among nutritional (restricted and unrestricted) and age (collection 1 and 2) groups; horizontally, among different categories of genes and viruses. Significant P showing negative or positive R-values, are represented in green or in red font. Likewise, percentages of correlations predominantly positives or negatives, are highlighted in green or red, respectively.

## REFERENCES

1. Klein A, Vaissière BE, Cane JH, Steffan-Dewenter I, Cunningham SA, et al. Importance of crop pollinators in changing landscapes for world crops. Proc Biol Sci. 2007;274(1608):303–13.

2. Kearns CA, Inouye DW, Waser NM Endangered mutualisms: The conservation of plant-pollinator interactions. Annu Rev Ecol Syst. 1998;29:83–112.

3. Vanengelsdorp D, Evans JD, Saegerman C, Mullin C, Haubruge E, et al. Colony collapse disorder: a descriptive study. PLoS One. 2009; 4 (8):e6481.

4. Neumann P, Carreck NL. Honey bee colony losses. J Apic Res. 2010;49(1):1–6.

5. Henry M, Béguin M, Requier F, Rollin O, Odoux JF, Aupinel P, Aptel J, Tchamitchian S, Decourtye A. A common pesticide decreases foraging success and survival in honey bees. Science. 2012;336(6079):348–50.

6. Naug D. Nutritional stress due to habitat loss may explain recent honeybee colony collapses. Biological Conservation. 2009;142:2369–72.

7. Dolezal A, Carrillo-Tripp J, Miller WA, Bonning BC, Toth AL. Intensively Cultivated Landscape and Varroa Mite Infestation Are Associated with Reduced Honey Bee Nutritional State. PLoS One. 2016;11(4):e0153531. doi: 10.1371/journal.pone.0153531.

8. Smart D, Pettis JS, Euliss N, Spivak MS. Land use in the Northern Great Plains region of the U.S. influences the survival and productivity of honey bee colonies. Agric Ecosyst Environ. 2016;230:139–49.

9. Amdam G, Hartfelder K, Norberg K, Hagen A, Omholt SW. Altered physiology in worker honey bees (Hymenoptera: Apidae) infested with the mite Varroa destructor (Acari: Varroidae): a factor in colony loss during overwintering? J Econ Entomol. 2004;97(3):741–7.

10. Guzmán-Novoa E, Eccles L, Calvete Y, Mcgowan J, Kelly PG, Correa-Benitez A. Varroa destructor is the main culprit for the death and reduced populations of overwintered honey bee (Apis mellifera) colonies in Ontario, Canada. Apidologie. 2010;41(4):443–50.

11. Martin S, Highfield AC, Brettell L, Villalobos EM, Budge GE, et al. Global honey bee viral landscape altered by a parasitic mite. Science. 2012;336(6086):1304–6.

12. Oldroyd B. What’s killing American honey bees? PLoS Biol. 2007;5(6):e168.

13. Ratnieks F, Carreck NL Clarity on honey bee collapse? Science. 2010; 327(5962):152–3.

14. Smart M, Pettis J, Rice N, Browning Z, Spivak M. Linking Measures of Colony and Individual Honey Bee Health to Survival among Apiaries Exposed to Varying Agricultural Land Use. PLoS One. 2016;11(3):e0152685. doi: 10.1371/journal.pone.0152685. PubMed Central PMCID: PMC27027871.

15. Decourtye A, Alaux C, Odoux J-F, Henry M, Vaissière BE, Le Conte Y. Chapter 16: Why Enhancement of Floral Resources in Agro-Ecosystems Benefit Honeybees and Beekeepers?. In: Grillo O VG, editor. Ecosystems Biodiversity: InTech; 2011. p. 371–88.

16. Haydak MH. Honey bee nutrition Annu Rev Entomol. 1970;15:143–56.

17. Herbert E. Honey bee nutrition. In: Graham J, editor. The Hive and the Honey Bee. Hamilton, Illinois: Dadant & Sons; 1992. p. 197–233.

18. Keller I, Fluri P, Imdorf I. Pollen nutrition and colony development in honey bees: part 1. Bee World. 2005;86(1):3–10.

19. Huang Z. Pollen nutrition affects honey bee stress resistance. Terr Arthropod Rev. 2012;5(175–189).

20. Manning R. Fatty acid composition of pollen and the effect of two dominant fatty acids (linoleic and linolenic) in pollen and flour diets on longevity and nutritional composition of honey bees (Apis mellifera) [PhD thesis]: Murdoch University, Western Australia, Australia; 2007.

21. Robinson GE. Regulation of division of labor in insect societies. Annu Rev Entomol. 1992;37:637–65.

22. Gordon D, Chu J, Lillie A, Tissot M, Pinter N. Variation in the transition from inside to outside work in the red harvester ant Pogonomyrmex barbatus. Insectes Sociaux. 2005;52(3):212–7.

23. Winston M. Biology of the Honey Bee. Cambridge, MA: Harvard University Press; 1987.

24. Rueppell O, Kaftanouglu O, Page RE. Honey bee (Apis mellifera) workers live longer in small than in large colonies. Exp Gerontol. 2009;44(6-7):447–52.. doi: 10.1016/j.exger.2009.04.003.

25. Goblirsch M, Huang ZY, Spivak M. Physiological and behavioral changes in honey bees (Apis mellifera) induced by Nosema ceranae infection. PLoS One. 2013;8(3):e58165. doi: 10.1371/journal.pone.0058165.

26. Schulz D, Huang ZY, Robinson GE Effects of colony food shortage on behavioral development in honey bees. Behav Ecol Sociobiol. 1998;42:295–303.

27. Nilsen K, Ihle KE, Frederick K, Fondrk MK, Smedal B, Hartfelder K, Amdam GV. Insulin-like peptide genes in honey bee fat body respond differently to manipulation of social behavioral physiology. J Exp Biol. 2011;214(9):1488–197.

28. Nelson C, Ihle KE, Fondrk MK, Page RE, Amdam GV. The gene vitellogenin has multiple coordinating effects on social organization. PLoS Biol. 2007; 5:e62.

29. Corona M, Velarde RA, Remolina S, Moran-Lauter A, Wang Y, et al. Vitellogenin, juvenile hormone, insulin signaling, and queen honey bee longevity. Proc Natl Acad Sci U S A. 2007;104(17):7128–33. doi: 17438290

30. Ament S, Corona M, Pollock HS, Robinson GE. Insulin signaling is involved in the regulation of worker division of labor in honey bee colonies. Proc Natl Acad Sci USA. 2008;105(11):4226–31.

31. Corona M, Libbrecht R, Wheeler DE. Molecular mechanisms of phenotypic plasticity in social insects. Curr Opin Insect Sci. 2016;13:55–60.

32. Raikhel A, Dhadialla TS Accumulation of yolk proteins in insect oocytes. Annu Rev Entomol. 1992;37(217-251).

33. Helvig C, Koener JF, Unnithan GC, Feyereisen R. CYP15A1, the cytochrome P450 that catalyzes epoxidation of methyl farnesoate to juvenile hormone III in cockroach corpora allata. Proc Natl Acad Sci U S A. 2004;101(12):4024–9.

34. Honeybee Genome Sequencing Consortium. Insights into social insects from the genome of the honeybee Apis mellifera. Nature. 2006; 443(7114):931–49.

35. Rutz W, Luscher M. The occurrence of vitellogenin in workers and queens of Apis mellifera and the possibility of its transmission to the queen. J Insect Physiol. 1974;20:897–909.

36. Bloch G, Wheeler DE, Robinson GE. Endocrine influences on the organization of insect societies. In: Pfaff D, editor. Hormones, Brain, and Behavior. 3. San Diego, CA.: Academic Press; 2002. p. 195–236.

37. Engels W. Occurrence and significance of vitellogenin in female castes of social Hymenoptera. Am Zool. 1974; 14:1229–37.

38. Rutz W, Gerig L, Wille H, Luscher M. The function of juvenile hormone in adult worker honeybees, Apis mellifera. J Insect Physiol. 1976;22:1485–91.

39. Hartfelder K, Engels W Social insect polymorphism: Hormonal regulation of plasticity in development and reproduction in the honeybee. Current Topics in Developmental Biology. 1998;40(45-77).

40. Pinto LZ, Bitondi MM, Simoes ZL Inhibition of vitellogenin synthesis in Apis mellifera workers by a juvenile hormone analogue, pyriproxyfen. J Insect Physiol. 2000;46:153–60.

41. Jaycox E, Skowronek W, Godfrey G. Behavioral changes in worker honey bees (Ais melljfera) induced by injections of a juvenile hormone mimic. Ann Entomol Soc Am. 1974;67:529–34.

42. Blanchard G, Orledge GM, Reynolds SE, Franks NR. Division of labour and seasonality in the ant Leptothorax albipennis: worker corpulence and its influence on behaviour. Anim Behav. 2000;59:723–38.

43. Toth A, Robinson GE. Worker nutrition and division of labour in honeybees. Anim behav. 2005;69:427–35.

44. Toth A, Varala K, Newman TC, Miguez FE, Hutchison SK, et al. Wasp gene expression supports an evolutionary link between maternal behavior and eusociality. Science. 2007;318(5849):441–4.

45. Ament SA, Wang Y, Robinson, GE. Nutritional regulation of division of labor in honey bees: toward a systems biology perspective. Wiley Interdiscip Rev Syst Biol Med. 2010;2(5):566–76.

46. Daugherty T, Toth AL, Robinson GE. Nutrition and division of labor: Effects on foraging and brain gene expression in the paper wasp Polistes metricus. Mol Ecol. 2011;20(24):5337–47.

47. Crailsheim K, Schneider LHW, Hrassnigg N, Buehlmann G, Brosch U, et al. Pollen consumption and utilization in worker honey bees (Apis mellifera carnica): dependence on individual age and function. J Insect Physiol. 1992;38(409-419).

48. Toth AL, Kantarovich S, Meisel AF, Robinson GE. Nutritional status influences socially regulated foraging ontogeny in honey bees. J Exp Biol. 2005;208:4641–9.

49. Dainat B, Evans JD, Chen YP, Gauthier L, Neumann P. Predictive markers of honey bee colony collapse. PLoS One. 2012;7(2):e32151.

50. Lavine M, Strand MR. Insect hemocytes and their role in immunity. Biochem Mol Biol. 2002;32(10):1295–309. PubMed Central PMCID: PMC12225920.

51. Elrod-Erickson M, Mishra S, Schneider D. Interactions between the cellular and humoral immune responses in Drosophila. Curr Biol. 2000;10(13): 781–4. PubMed Central PMCID: PMC10898983.

52. Evans J, Aronstein K, Chen YP, Hetru C, Imler JL, et al. . Immune pathways and defence mechanisms in honey bees Apis mellifera. Insect Mol Biol. 2006;15:645–56.

53. Lemaitre B. The road to Toll. Nat Rev Immunol. 2004;4(7):521–7.

54. Casteels P, Ampe C, Riviere L, Van Damme J, Elicone C et al. Isolation and characterization of abaecin, a major antibacterial response peptide in the honeybee (Apis mellifera). Eur J Biochem. 1990;187(2):381–6.

55. Casteels-Josson K, Zhang W, Capaci T, Casteels P, Tempst P. Acute transcriptional response of the honeybee peptide-antibiotics gene repertoire and required post-translational conversion of the precursor structures. J Biol Chem. 1994;269(46):28569–75.

56. Casteels P, Ampe C, Jacobs F, Tempst P. Functional and chemical characterization of Hymenoptaecin, an antibacterial polypeptide that is infection-inducible in the honeybee (Apis mellifera). J Biol Chem. 1993; 268(10):7044–54. PubMed Central PMCID: PMC8463238.

57. Strand M, Pech LL. Immunological basis for compatibility in parasitoid-host relationships. Annu Rev Entomol. 1995;40:31–56.

58. Dziarski R, Gupta D. The peptidoglycan recognition proteins (PGRPs). Genome Biol. 2006;7(8):232. PubMed Central PMCID: PMC16930467.

59. Kocks C, Cho JH, Nehme N, Ulvila J, Pearson AM, et al. Eater, a transmembrane protein mediating phagocytosis of bacterial pathogens in Drosophila. Cell. 2005;123(2):335–46. PubMed Central PMCID: PMC16239149.

60. Söderhäll K, Cerenius L. Role of the prophenoloxidase-activating system in invertebrate immunity. Curr Opin Immunol. 1998;10(1):23–8.

61. González-Santoyo I, Córdoba-Aguilar, A. Phenoloxidase: a key component of the insect immune system. Entomol Exp Appl. 2012;142:1–16.

62. Alaux C, Ducloz F, Crauser D, Le Conte Y. Diet effects on honeybee immunocompetence. Biol Lett. 2010;6(4):562–5.

63. Alaux C, Dantec C, Parrinello H, Le Conte Y. Nutrigenomics in honey bees: digital gene expression analysis of pollen’s nutritive effects on healthy and varroa-parasitized bees. BMC Genomics. 2011;12:496.

64. Corby-Harris V, Jones BM, Walton A, Schwan MR, Anderson KE. Transcriptional markers of sub-optimal nutrition in developing Apis mellifera nurse workers.. BMC Genomics. 2014;15:(134). doi: 10.1186/1471-2164-15-134.

65. DeGrandi-Hoffman G, Chen Y, Huang E, Huang MH. The effect of diet on protein concentration, hypopharyngeal gland development and virus load in worker honey bees (Apis mellifera L.). J Insect Physiol. 2010;56(9):1184–91.

66. Di Pasquale G, Salignon M, Le Conte Y, Belzunces LP, Decourtye A, Kretzschmar A, Suchail S, Brunet JL, Alaux C. Influence of pollen nutrition on honey bee health: do pollen quality and diversity matter? PLoS One. 2013;8(8):e72016.

67. Tritschler M, Vollmann JJ, Yañez O, Chejanovsky N, Crailsheim K, Neumann P. Protein nutrition governs within-host race of honey bee pathogens. Sci Rep. 2017;7(1):14988. doi: 10.1038/s41598-017-15358-w.

68. Basualdo M, Barragán, S, Antúnez. Bee bread increases honeybee haemolymph protein and promote better survival despite of causing higher Nosema ceranae abundance in honeybees. Environ Microbiol Rep. 2014; 6:396–400. doi: https://doi.org/10.1111/1758-2229.12169.

69. Fewell J, Winston ML Colony state and regulation of pollen foraging in the honey bee, Apis mellifera L. Behav Ecol Sociobiol. 1992;30:387–93.

70. Dreller C, Page RE, Fondrk MK. Regulation of pollen foraging in honeybee colonies: effects of young brood, stored pollen, and empty space. Behav Ecol Sociobiol. 1999;45:227–33.

71. Pankiw T, Page RE, Fondrk MK. Brood pheromone stimulates pollen foraging in honey bees (Apis mellifera). Behav Ecol Sociobiol. 1998;44:193–8.

72. Page R, Scheiner R, Erber J, Amdam GV. The Development and Evolution of Division of Labor and Foraging Specialization in a Social Insect (Apis mellifera L.). Curr Top Dev Biol. 2006;74:253–86.

73. Keller I, Fluri P, Imdorf I. Pollen nutrition and colony development in honey bees: part II. Bee World 2005;86:27–34.

74. Perry C, Søvik E, Myerscough MR, Barron AB. Rapid behavioral maturation accelerates failure of stressed honey bee colonies. Proc Natl Acad Sci U S A. 2015;Feb 9.

75. Guzmán–Novoa E, Page RE, Gary NE. Behavioral and life history components of division of labors in honey bees. Behav Ecol Sobiobiol. 1994;34(6):409–17.

76. Rueppell O, Bachelier C, Fondrk MK, Page RE Jr. Regulation of life history determines lifespan of worker honey bees (Apismellifera L.). Exp Gerontol 2007;42(10):1020–32.

77. Piulachs M, Guidugli KR, Barchuk AR, Cruz J, Simões ZL, et al. The vitellogenin of the honey bee, Apis mellifera: structural analysis of the cDNA and expression studies. Insect Biochem Mol Biol. 2003;33(4):459–65.

78. Cremonez T, De Jong D, Bitondi MM. Quantification of Hemolymph Proteins as a Fast Method for Testing Protein Diets for Honey Bees (Hymenoptera: Apidae). J Econ Entomol 1998;91(6):1284–9.

79. Di Pasquale G, Alaux C, Le Conte Y, Odoux JF, Pioz M, Vaissière BE, Belzunces LP, Decourtye A. Variations in the Availability of Pollen Resources Affect Honey Bee Health. PLoS One. 2016;11(9):e0162818.

80. Lourenço A, Martins JR, Torres FAS, Mackert A, Aguiar LR, Hartfelder K, Bitondi MMG, Simões ZLP. Immunosenescence in honey bees (Apis mellifera L.) is caused by intrinsic senescence and behavioral physiology. Exp Gerontol. 2019;119:174–83.

81. Fluri P, Wille H, Gerig L, Lüscher M Juvenile hormone, vitellogenin and haemocyte composition in winter worker honeybees (Apis mellifera). Experimentia. 1977; 33:1240–1.

82. Seehuus SC, Norberg K, Gimsa U, Krekling T, Amdam GV. Reproductive protein protects functionally sterile honey bee workers from oxidative stress. Proc Natl Acad Sci U S A. 2006;03(4):962–7.

83. Drapeau M, Albert S, Kucharski R, Prusko C, Maleszka R. Evolution of the Yellow/Major Royal Jelly Protein family and the emergence of social behavior in honey bees. Genome Res. 2006;16(11):1385–94.

84. Niu D, Zheng H, Corona M, Lu Y, Chen X, Cao L, Sohr A, Hu F. Transcriptome comparison between inactivated and activated ovaries of the honey bee Apis mellifera L Insect Mol Biol. 2014;23(5):668–81. doi: 10.1111/imb.12114. PubMed Central PMCID: PMC25039886.

85. Ji T, Liu Z, Shen J, Shen F, Liang Q, Wu L, Chen G, Corona M. Proteomics analysis reveals protein expression differences for hypopharyngeal gland activity in the honeybee, Apis mellifera carnica Pollmann. BMC Genomics. 2014;15:665.

86. Crailsheim K. The flow of jelly within a honeybee colony. Journal of Comparative Physiology B 1992;162(8):681–9.

87. Camazine S. The regulation of pollen foraging by honey bees: how foragers assess the colony’s need for pollen. Behav Ecol Sociobiol. 1993;32:265–72.

88. Camazine S, Crailsheim K, Hrassnigg N, Robinson GE, Leonhard B, et al. Protein trophallaxis and the regulation of pollen foraging by honey bees (Apis mellifera L.). Apidologie. 1998;29:113–26.

89. Bomtorin A, Mackert A, Rosa GC, Moda LM, Martins JR et al. Juvenile hormone biosynthesis gene expression in the corpora allata of honey bee (Apis mellifera L.) female castes. PLoS One. 2014;9(1):e86923.

90. Tian L, Ji BZ, Liu SW, He CL, Jin F, Gao J, Stanley D, Li S. JH biosynthesis by reproductive tissues and corpora allata in adult longhorned beetles, Apriona germari. Arch Insect Biochem Physiol. 2010;75(4):275–86.

91. Borovsky D, Carlson DA, Ujvary I, Prestwich GD. Biosynthesis of (10R)-Juvenile hormone III from farnesoic acid by Aedes aegypti ovary. Arch Insect Biochem Physiol 1994;2:75–90.

92. Bellés X, Martín D, Piulachs MD. The mevalonate pathway and the synthesis of juvenile hormone in insects. Annu Rev Entomol. 2005;50:181–99.

93. Sullivan J, Fahrbach SE, Harrison JF, Capaldi EA, Fewell JH, et al. Juvenile hormone and division of labor in honey bee colonies: effects of allatectomy on flight behavior and metabolism. J Exp Biol. 2003;206(Pt 13):2287–96.

94. Moret Y, Schmid-Hempel P. Survival for immunity: the price of immune system activation for bumblebee workers. Science. 2000;290(5494):1166–8. PubMed Central PMCID: PMC11073456.

95. Di Prisco G, Cavaliere V, Annoscia D, Varricchio P, Caprio E, Nazzi F, Gargiulo G, Pennacchio F. Neonicotinoid clothianidin adversely affects insect immunity and promotes replication of a viral pathogen in honey bees. Proc Natl Acad Sci U S A 2013;110(46):18466–71. doi: 10.1073/pnas.1314923110.

96. Huang Z, Robinson GE. Honeybee colony integration: worker-worker interactions mediate hormonally regulated plasticity in division of labor. Proc Natl Acad Sci U S A. 1992;89(24):11726–9.

97. Rutz W, Gerig L, Wille H, Luscher, M. A bioassay for juvenile hormone (JH) effects of insect growth regulators (IGR) on adult worker honeybees. Bull de la Soc Entomol Suisse. 1974;47:307–13.

98. Amdam G, Aase AL, Seehuus SC, Kim Fondrk M, Norberg K, Hartfelder K. Social reversal of immunosenescence in honey bee workers. Exp Gerontol. 2005;40(12):939–47.

99. Amdam G, Simões ZL, Hagen A, Norberg K, Schrøder K, et al. Hormonal control of the yolk precursor vitellogenin regulates immune function and longevity in honeybees. Exp Gerontol. 2004;39(5):767–73.

100. Schmid M, Brockmann A, Pirk CW, Stanley DW, Tautz J. Adult honeybees (Apis mellifera L.) abandon hemocytic, but not phenoloxidase-based immunity. J Insect Physiol. 2008;54(2):439–44.

101. Steinmann N, Corona M, Neumann P, Dainat B. Overwintering Is Associated with Reduced Expression of Immune Genes and Higher Susceptibility to Virus Infection in Honey Bees. PLoS One. 2015;10(6):e0129956.

102. Bull J, Ryabov EV, Prince G, Mead A, Zhang C, Baxter LA, Pell JK, Osborne JL, Chandler D. A strong immune response in young adult honeybees masks their increased susceptibility to infection compared to older bees. PLoS Pathog. 2012;8(12):e1003083.

103. Corona M, Hughes KA, Weaver DB, Robinson GE. Gene expression patterns associated with queen honey bee longevity. Mech Ageing Dev. 2005;126(11):1230–8.

104. Wilson-Rich N, Dres ST, Starks PT. The ontogeny of immunity: development of innate immune strength in the honey bee (Apis mellifera). J Insect Physiol. 2008;54(10-11):1392–9.

105. Laughton A, Boots M, Siva-Jothy MT. The ontogeny of immunity in the honey bee, Apis mellifera L. following an immune challenge. J Insect Physiol. 2011;57(7):1023–32.

106. Dudzic J, Kondo S, Ueda R, Bergman CM, Lemaitre B. Drosophila innate immunity: regional and functional specialization of prophenoloxidases. BMC Biol. 2015;13:81.

107. Christophides G, Zdobnov E, Barillas-Mury C, Birney E, Blandin S, et al. Immunity-related genes and gene families in Anopheles gambiae. Science. 2002;298:159–65.

108. Lourenco A, Zufelato MS, Bitondi MMG, Simoes ZLP. Molecular characterization of a cDNA coding prophenoloxidase and its expression in Apis mellifera. Insect Biochem Mol Biol. 2005;35:541–52.

109. Kanost M, Gorman MJ. Phenoloxidases in insect immunity. In: Beckage N, editor. Insect Immunology. San Diego, CA, USA: Academic Press; 2008. p. 69–96.

110. Shelby K, Popham HJ. Plasma phenoloxidase of the larval tobacco budworm, Heliothis virescens, is virucidal. J Insect Sci. 2006; 6:1–12.

111. Beck M, Strand MR. A novel polydnavirus protein inhibits the insect prophenoloxidase activation pathway. Proc Natl Acad Sci U S A. 2007;104(49):19267–72.

112. Yang X, Cox-Foster DL. Impact of an ectoparasite on the immunity and pathology of an invertebrate: evidence for host immunosuppression and viral amplification. Proc Natl Acad Sci U S A. 2005;102(21):7470–5.

113. Ryabov E, Fannon JM, Moore JD, Wood GR, Evans DJ. The Iflaviruses Sacbrood virus and Deformed wing virus evoke different transcriptional responses in the honeybee which may facilitate their horizontal or vertical transmission. PeerJ 2016;4:e1591.

114. Giray T, Robinson GE. Effects of intracolony variability in behavioral development on plasticity of division of labor in honey bee colonies. Behav Ecol Sociobiol. 1994;35:13–20.

115. Gregorc A, Evans JD, Scharf M, Ellis JD. Gene expression in honey bee (Apismellifera) larvae exposed to pesticides and Varroa mites (Varroa destructor). J Insect Physiol. 2012;58(8):1042–9.

116. Livak KJ, Schmittgen TD. Analysis of relative gene expression data using real-time quantitative PCR and the 2-[Delta][Delta] CT method. methods. 2001;25(4):402–8.

117. Rychlik W. OLIGO 7 primer analysis software. METHODS IN MOLECULAR BIOLOGY-CLIFTON THEN TOTOWA-. 2007;402:35.

118. Gross J, Ligges U. Nortest: Tests for Normality. R package version 1.0-4. 2015.

119. Fox J, Weisberg S. An {R} Companion to Applied Regression. Second Edition ed. Thousand Oaks CA: Sage; 2011.

120. RStudio Team. RStudio: Integrated Development for R. Boston, MA: RStudio Inc; 2016.

121. de Mendiburu F. Agricolae: Statistical Procedures for Agricultural Research. R package version 1.2–4. 2016.

122. Oksanen J, Blanchet FG, Friendly M, Kindt R, Legendre P, et al. vegan: Community Ecology Package. R package version 2.4-3. 2017.

